# Energy-dependent transport at dural lymphatic vessels is necessary for Aβ brain clearance in Alzheimer’s disease

**DOI:** 10.1101/427617

**Authors:** Liudmila Romanova, Heidi Phillips, Gregory S. Calip, Kyle Hauser, Daniel A. Peterson, Orly Lazarov, Daniel Predescu, Sanda Predescu, Julie Schneider, Jeff Kordower, Eric Hansen, Cornelius H. Lam, Christopher G. Janson

## Abstract

Viewed as an imbalance between production and clearance of toxic Aβ peptides, Alzheimer’s disease is a candidate for therapies to augment brain waste removal. Prior work has shown that Aβ accumulates in meninges with aging as a byproduct of normal brain activity, in parallel with build-up of Aβ oligomers in neurons, blood vessels, and interstitial fluid. Using the TgF344-AD rat model of Alzheimer’s disease, we now report that dural lymphatic vessels specifically accumulate neurotoxic pyroglutamate amyloid beta (pE3-Aβ) with aging. Notably, accelerated amyloidosis is observed in meninges after ligation of cervical lymphatics, together with significantly increased pE3-Aβ and Aβ42 deposition in upstream brain regions implicated in Alzheimer’s disease. Blockade of lymphatic clearance is not sufficiently compensated by other efflux pathways, suggesting a necessary role of Aβ clearance at the level of lymphatics. We further report that dural lymphatic cells actively clear Aβ via energy-dependent mechanisms, and lymphatic Aβ transport is significantly impaired both in normal aging and in Alzheimer’s disease. Dural lymphatic cells isolated from the TgF344-AD rat show ultrastructural abnormalities in mitochondria and abnormal cytoplasmic inclusions, with a distinct transcriptional profile implicating failure of energy-dependent transport. Finally, using human meninges treated with FocusDeep tissue clearing, we demonstrate using whole mount panoramic imaging that dural lymphatic vessels comprise a structurally diverse intracranial vascular network that accumulates pE3-Aβ with aging, similar to the rat model. We conclude that intracranial meningeal and extracranial cervical lymphatic vessels are targets for Alzheimer’s disease therapies focused on improving amyloid clearance.

**One Sentence Summary:** Lymphatic vessels remove Aβ from the brain via energy-dependent active transport mechanisms, and blockage of extracranial lymphatic drainage is sufficient to cause significant acceleration of intracranial Alzheimer’s Aβ pathology in both meninges and brain.

## Introduction

The brain lacks an intrinsic lymphatic vasculature, which in other organs is critical for clearance of excess fluid and protein from the interstitial space as well as immune surveillance. Without a functioning lymphatic system, interstitial tissues of the body tend to become edematous and polluted. Yet, despite the absence of lymphatics in the brain, it paradoxically maintains proper fluid and solute homeostasis. Part of the explanation for this incongruity involves a system of tightly regulated fluid and solute transport through the blood-brain barrier at the level of cerebral capillaries. In addition, paravascular drainage routes (*1*) connect brain interstitial fluid deep inside the brain with the subarachnoid space and cerebrospinal fluid (CSF). Known for many years as “pre-lymphatic” drainage (*2*), these paravascular channels were recently termed “glymphatics” due to a putative role of glia in promoting bulk fluid flow (*3*). Excess brain fluids and solutes are ultimately drained out through the meninges, skin-like protective membranes surrounding the brain, and their integrated venous sinuses which function as a terminal hub for waste clearance.

An important unresolved question in physiology relates to how the meninges work together with the cerebral vasculature to maintain brain health. Far from being an inert protective sac, the meninges are a complex and dynamic organ with multiple vascular beds and diverse resident cells. The outer meningeal layer of dura contains a rich microvasculature consisting of arteries, veins, and lymphatics as well as large amounts of hydrated collagen and other connective tissue which contacts underlying arachnoid membranes. Relatively few studies have examined this vascular network and its cellular milieu in detail. Like the blood-brain barrier inside the brain, the membranes of the meninges provide an interface between discrete fluid compartments. While the arterial blood supply into the brain and dura are distinct, diverging at the internal and external carotid arteries, all of the brain’s exiting venous blood and much of its daily surplus of cerebrospinal fluid (CSF) flows out through dural venous sinuses. Dural lymphatic vessels are known to abut these central venous structures in rodents and other mammals, and exit through the skull base and cranium. Although they are presumed to have a role in clearance of brain fluids and solutes, the architecture and physiological significance of dural lymphatics in humans has been unclear.

The meningeal lymphatic system was first discovered in 1787 by Mascagni (*4*) who used mercuric injections to map its gross anatomy in humans post-mortem **(fig. S1)**. A century later, the first experiments demonstrating lymphatic drainage from the subarachnoid space were conducted circa 1870. Using vital dyes, investigators showed that in addition to outflow through arachnoid granulations, CSF and solutes exit the skull base along cranial nerves, with passage to nasal and cervical lymphatics (*5*). The unique role of meninges in CSF and solute drainage was demonstrated by Weed in 1914, who was also aware of a lymphatic contribution (*6*). Impaired dural lymphatic drainage as an underlying cause of brain pathology was proposed in 1943 by Speransky, who posited a direct connection between the subarachnoid space and the lymphatic system of the body (*7*). Lymphatic clearance of brain fluids through meninges was revisited by Foldi in the 1960s, in his studies on “lymphogenic encephalopathy” (8,*9*). Drainage from brain to extracranial lymphatics was confirmed in multiple mammalian species over subsequent decades (*10-12*), with a continued emphasis on skull base pathways rather than along the cranial vault. Despite this extensive body of work, the role of meningeal lymphatics in human disease has remained speculative, with hydrocephalus as the main implicated pathology. Outside of the neurosurgical literature, these early anatomical discoveries were not widely appreciated. In fact, one of the 20^th^ century leaders in brain barrier studies candidly admitted that his lab was initially “unaware of previous work on CSF connections with the lymphatic system” as they began tracer experiments in the 1970s *(13).*

Before the discovery of Aβ peptide, it was shown that ligation of cervical lymphatic vessels in animals led to cognitive impairment (*14*), but a link to meningeal lymphatics or Alzheimer’s disease was never considered. It was observed in those early ligation studies that there was dilation of perivascular spaces in the brain, with proteinaceous material evident on electron microscopy. Years later, a seminal study showed that injection of fluorescently labeled Aβ to the rat brain is cleared by perivascular drainage routes to both meninges and deep cervical lymph nodes (*15*). It was also reported that significant Aβ and pyroglutamate Aβ (pE3-Aβ) accumulation occurs in cervical and axillary lymph glands of aged Alzheimer’s disease mice, which suggests that endogenous brain Aβ may be cleared through lymphatic drainage (*16*). Subsequently, transgenic mice expressing lymphatic-specific markers were used to visualize the structure of lymphatic microvasculature in mouse meninges; targeted blockade of either cervical lymphatics (*17*) or meningeal lymphatics (*18*) in those models prevented drainage of peptide tracers, confirming prior results that lymphatic vessels are involved in brain solute clearance. Recently, photochemical ablation of intracranial lymphatics in a mouse model of Alzheimer’s disease was associated with increased hippocampal Aβ and cognitive impairment; in animals with impaired meningeal lymphatics, investigators also reported greater retention of labeled Aβ in meninges (*19*). Altogether, these studies suggest a possible link between meningeal and cervical lymphatic function and Aβ peptide clearance, in addition to the accepted role of cerebral blood vessels.

Correlative studies have consistently shown that Aβ accumulates in human meninges with aging *(20-23),* but whether amyloid deposition in this location is causative or incidental to disease progression has been an important open question. Given that Alzheimer’s disease is correlated with deficient clearance of Aβ peptide (*24*) and the meninges are a terminal hub for brain waste clearance, our lab’s central hypothesis has been that impaired Aβ clearance through meninges influences Alzheimer’s disease progression. More specifically, in this study we hypothesized that impaired lymphatic clearance of Aβ accelerates the normal aging process and potentiates Alzheimer’s disease brain pathology. We tested the hypothesis of a causal relationship between solute clearance and Alzheimer’s disease progression by measuring different species of Aβ in hippocampus and cortical areas before and after cervical lymphatic ligation. Here we report that intracranial and extracranial lymphatic vessels are functionally linked through the meninges, and are necessary for brain clearance of highly toxic pE3-Aβ, a known key player in Alzheimer’s disease pathogenesis.

It is generally accepted in the lymphatic biology field that paracellular or passive transport is the primary means of lymphatic uptake of fluid and solutes *(25),* and thus our additional finding that meningeal lymphatics use active transport to clear Aβ was surprising and unexpected. Although lymphatics appear to clear a variety of Aβ isoforms, the preferred uptake of pE3-Aβ by meningeal lymphatics was intriguing, given its ability to seed fibrillar perineuronal deposits as the so-called “hatchet man” of Alzheimer’s disease *(26).* Our data suggest that lymphatic outflow tracts are necessary for clearance of toxic pE3-Aβ from the brain, and that differential transport of Aβ species occurs in specific endothelial vessels. In this paradigm, pE3-Aβ clearance flows from neurons to interstitial fluid to perivascular spaces to meninges to lymphatic capillaries and collecting vessels. It is known that familial Aβ isoforms cross endothelia to variable degrees *(27)* and that cerebral amyloid angiopathy predominantly involves the Aβ40 isoform *(28)*. But until now, differential microvascular uptake of post-translationally modified Aβ species has not been described, which opens up new opportunities for targeted drug discovery.

For experimental studies, we used the Tg-F344-AD rat model that exhibits key hallmarks of human Alzheimer’s disease such as Aβ and tau pathology, synaptic loss, and age-dependent cognitive impairment *(29).* Of major significance to our study, this rat model has significant and early accumulation of pE3-Aβ, which is present in human disease but generally lacking in mouse models *(30)*. In order to distinguish effects of age, disease, and length of ligation, we microsurgically treated young and old TgF344-AD rats and used a generalized linear model to assess these independent variables. Primary outcome variables for *in vivo* studies were biochemical and immunohistochemical assays of amyloid burden in the meninges and brain. In addition, we isolated pure populations of lymphatic cells for *in vitro* mechanistic studies and RNAseq expression analysis. Finally, using freshly harvested human meninges, we verified that observations in the Tg-F344-AD rat were consistent with human Alzheimer’s disease pathology, and mapped the existence of a dural lymphatic network in humans using tissue clearing techniques with full-thickness whole mounts.

Our data support the following novel findings: (1) in the dura mater, lymphatic capillaries and collecting ducts uptake neurotoxic pE3-Aβ; (2) restriction of extracranial lymphatic drainage in the neck is sufficient to cause accumulation of pE3-Aβ and Aβ42 in meninges, skull, and cognitively important brain regions; (3) consistent with a clearance function, meninges and cervical lymph nodes accumulate pE3-Aβ at early disease stages, prior to advanced neuronal pathology; (4) dural lymphatic cells use energy-dependent mechanisms of Aβ uptake that involve both endocytosis and multi-drug transporters, which can be pharmacologically targeted; (5) dural lymphatic cells derived from TgF344-AD rats have severely impaired Aβ transport, along with distinctive ultrastructural changes in aging which suggest mitochondrial dysfunction; (6) dural lymphatic cells derived from TgF344-AD rats show significant differences in gene expression compared to wild-type, consistent with deficiencies in energy-dependent transport; (7) human meninges contain a similar network of dural lymphatic vessels which are well-visualized following tissue clearing, and which specifically accumulate pE3-Aβ in the setting of Alzheimer’s disease. These results suggest that the meninges and lymphatic system of the head and neck represent a target for early interventions in Alzheimer’s disease prevention and therapy and warrant further investigations.

## Results

### Rat dural lymphatics have diverse morphologies & complex relationships to other vascular beds

Most recent studies on meningeal lymphatics have been conducted in mice. As with mouse meninges, the dura mater of rats is diaphanous, similar in appearance to the delicate arachnoid layer in humans. However, in addition to having a more complex vasculature, rat meninges are thicker and can be stripped from the inner table of the skull in the same manner as with human neurosurgical procedures. Thus, unfixed meninges are readily collected for both immunohistochemical staining and biochemical studies. Our lab has developed tissue staining, clearing, and mounting techniques in order to view lymphatic capillaries and collecting ducts in whole meninges up to 500μm in the Z-plane, using the TgF344-AD model of Alzheimer’s disease. Using fast resonance mosaic confocal scanning and standard confocal, we serially imaged wild-type and Alzheimer’s rat meninges to assess different vascular beds, which demonstrate discrete lymphatic networks closely associated with meningeal arteries and veins **[Fig.1]**. The distribution of these vascular beds suggests that there are at least three distinct lymphatic drainage pathways from the brain: along central venous sinuses, along the middle meningeal artery, and along olfactory tracts **(fig. S2)**. Higher magnification views using aqueous clearing agents **[Fig.2]** demonstrate fine lymphatic vessels which accompany both meningeal venous sinuses and arterial beds.

**Fig. 1.**
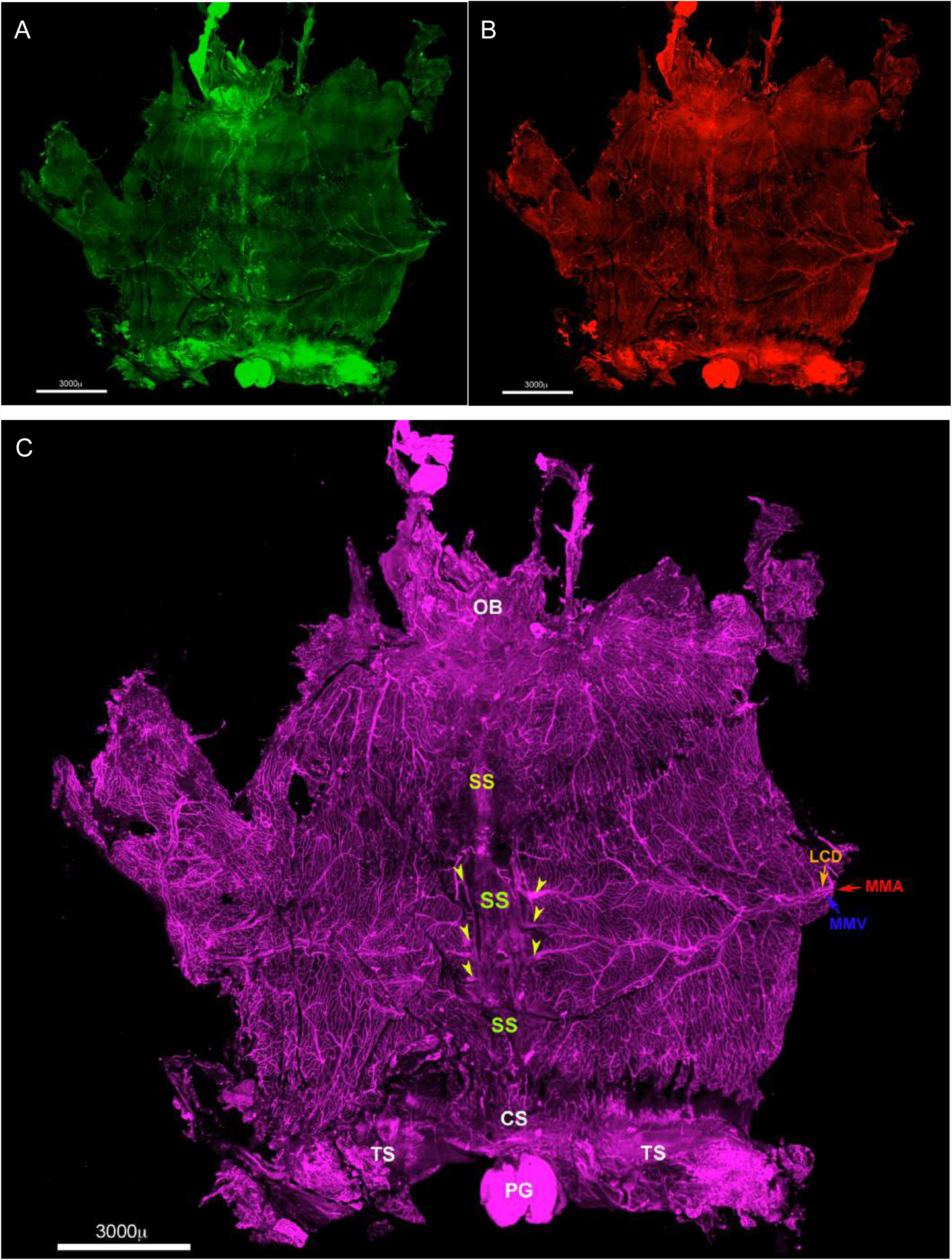
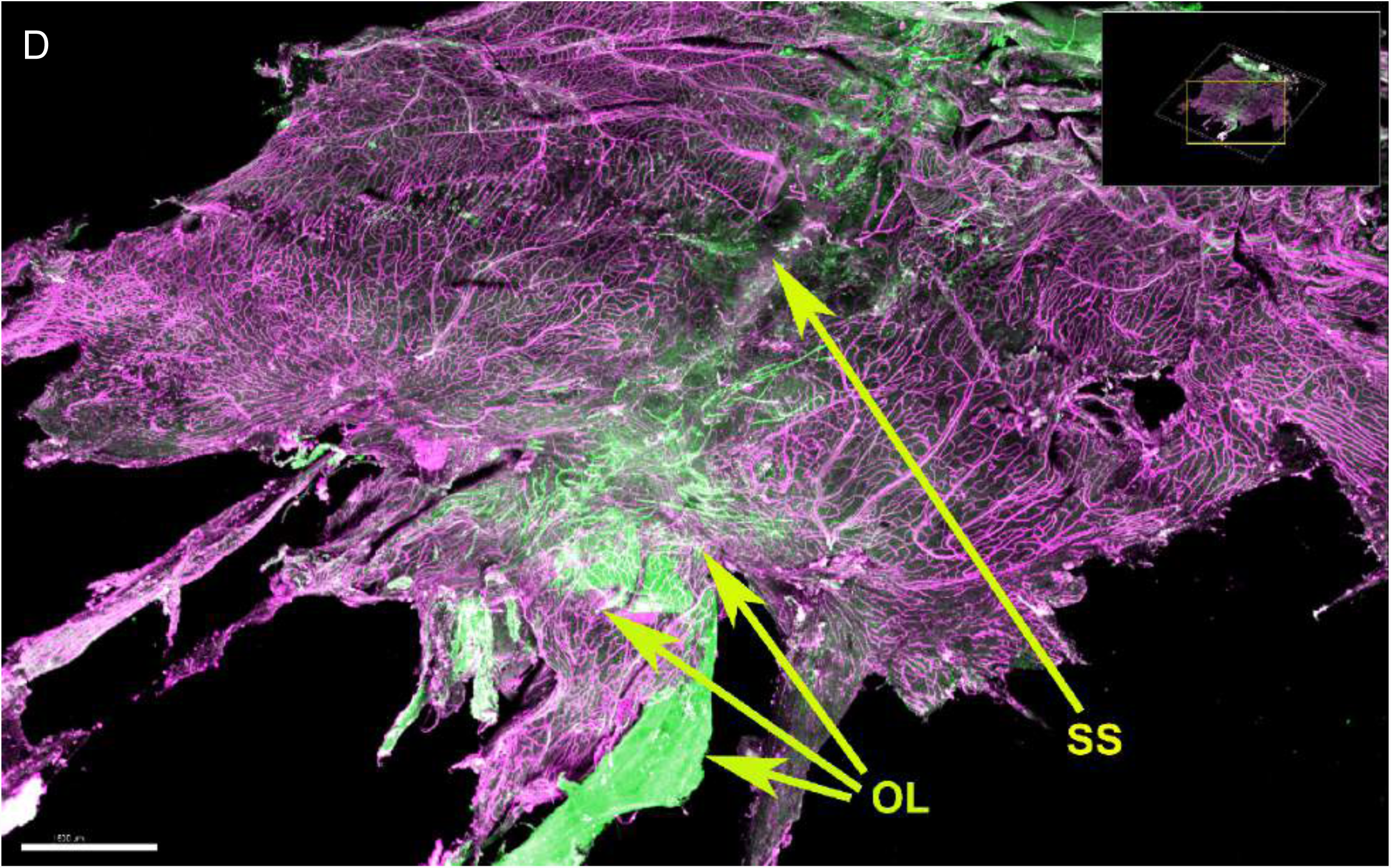
Rat meninges are richly vascularized with a diverse network of veins, arteries, and lymphatics. Fig. 1 Legend. Whole mount meninges, staining to Pdpn (A) and Lyve1 (B), showing networks of lymphatics concentrated along venous sinuses, olfactory region, and meningeal arteries. The same specimen (C) is stained to generic vascular marker CD31 (C), showing olfactory bulb (OB), superior sagittal sinus (SS), transverse sinus (TS), pineal gland (PG), middle meningeal artery and vein (MMA, MMV), and lymphatic collecting ducts (LCD). Yellow arrows denote large draining veins into the central venous sinus, which anastomose with laterally directed arteries. Anterior view (D) of olfactory lymphatics (OL) demonstrates strong staining to Pdpn (green) alongside other vasculature.

**Fig. 2.**
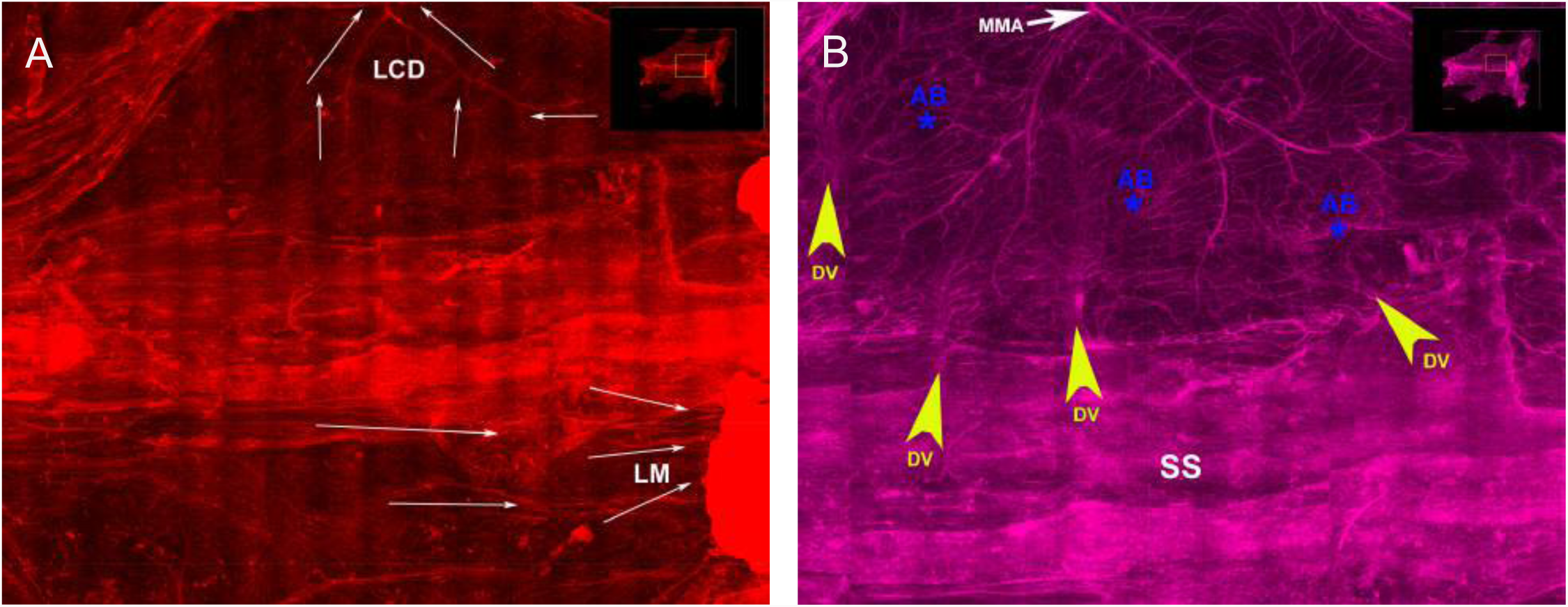
Cleared meninges demonstrate microvascular drainage routes around sagittal sinus. Fig. 2 Legend. Whole meninges cleared with FocusDeep, showing staining to Lyve1 (A) and to CD31 (B). Networks of lymphatics run medially along the sagittal venous sinus, and laterally along meningeal arteries. White arrows indicate the presumed direction of flow through lymphatic collecting ducts (LCD) and lymphatic microvasculature (LM). Yellow arrows indicate large draining veins and blue asterisks indicate anastomotic beds between arteries and veins.

### Aβ species accumulate with aging in meninges, which is potentiated by lymphatic ligation

TgF344-AD animals uniformly accumulate substantial quantities of Aβ species in meninges with aging, as assessed by immunohistochemistry and biochemical assays. Aged wild-type animals accumulate smaller amounts of amyloid with aging. We found that meningeal Aβ deposition is potentiated following cervical lymphatic ligation, confirming a direct link between intracranial and extracranial lymphatic drainage routes, as originally proposed by Speransky over 75 years ago. Immunohistochemistry of whole meninges from aged animals showed differences in Aβ signal between ligated and non-ligated wild-type and TgF344-AD animals **[Fig.3]**, with ligated animals having more diffuse Aβ signal than non-ligated, and TgF344-AD animals having the highest amyloid burden. Comparing ligated and non-ligated TgF344-AD animals, there were quantitative differences in the intensity of Aβ signal, evident in multiple areas of dura including the lateral meninges bordering the middle meningeal artery **[Fig.4]**. The level of amyloid deposition and baseline autofluorescence along the central venous sinuses was uniformly high. These results suggest that cervical lymphatic ligation causes a relative blockade of amyloid drainage, mediated in part through lateral meningeal lymphatics and paravascular flow surrounding the middle meningeal artery. In selected TgF344-AD animals, we examined the demineralized calvarium for the presence of endothelial channels together with Aβ signal. We found numerous diploic channels and resident cells containing pE3-Aβ and pan-Aβ signal **(fig. S3)**. This suggests that vascular routes at the vertex also may be involved in Aβ clearance.

**Fig. 3.**
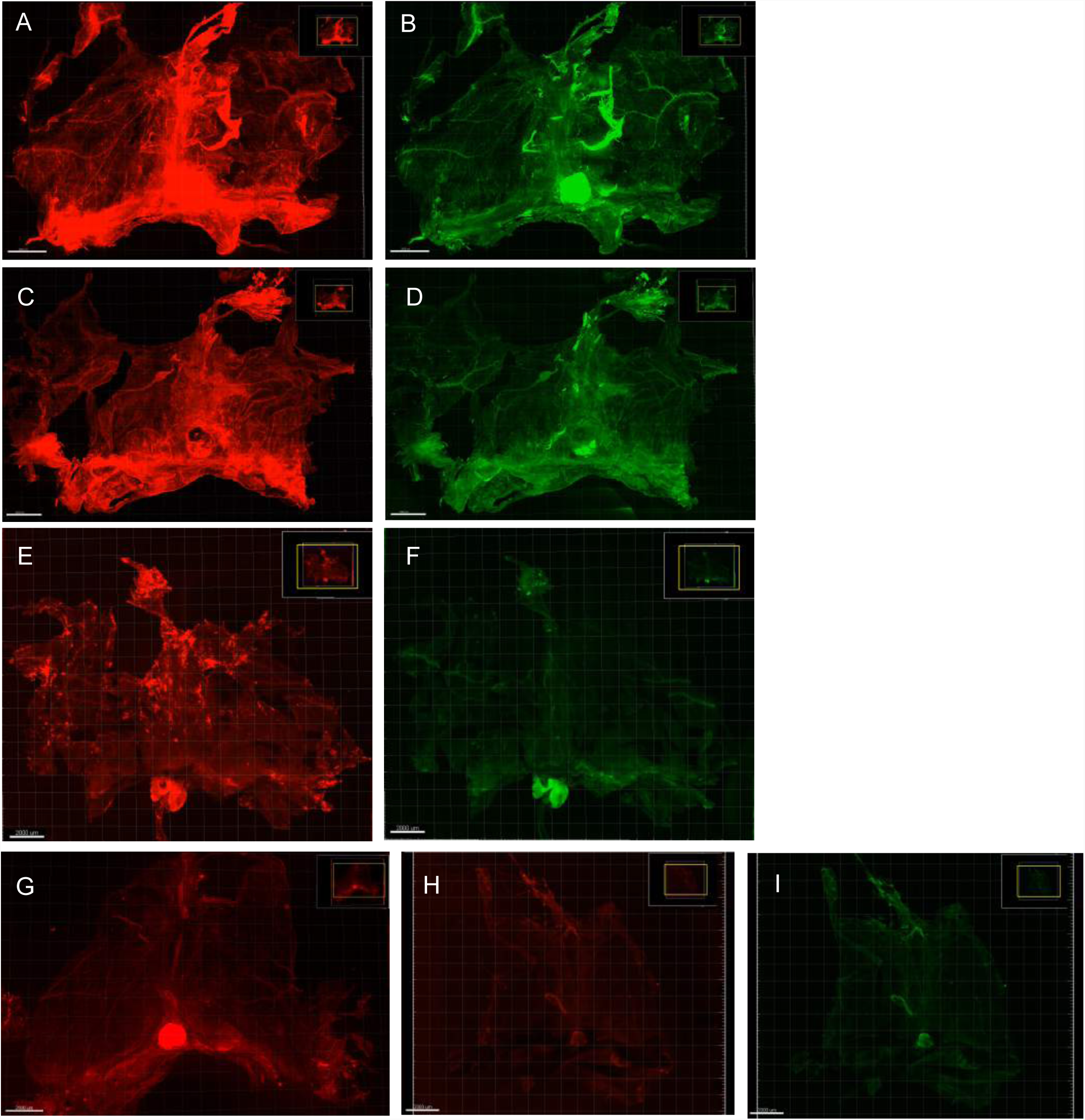
Effects of cervical ligation and aging on meningeal amyloidosis. Fig. 3 Legend. Whole meninges from TgF344-AD and wild-type animals, showing effects of ligation and aging on amyloid accumulation. 18 month old TgF344-AD rat following ligation for 3 months, showing (A) strong pan-Aβ signal (B) and pE3-Aβ signal. Age-matched 18 month old TgF344-AD without ligation, showing (C) pan-Aβ signal and (D) pE3-Aβ signal. The pan-Aβ signal along venous sinuses appears greater in the ligated animal, while pE3-Aβ signal appears similar. Images were acquired under identical confocal settings. 21 month old wild-type rat following cervical lymphatic ligation for 6 months shows relative increase in diffuse pan-Aβ signal (E) and pE3-Aβ signal (F), compared to 18 month wild-type animal without ligation (G), suggesting that ligation independently potentiates buildup of meningeal amyloid. Age-matched wild-type negative controls (H,I) without primary antibody show background autofluorescence on red and green channels.

**Fig. 4.**
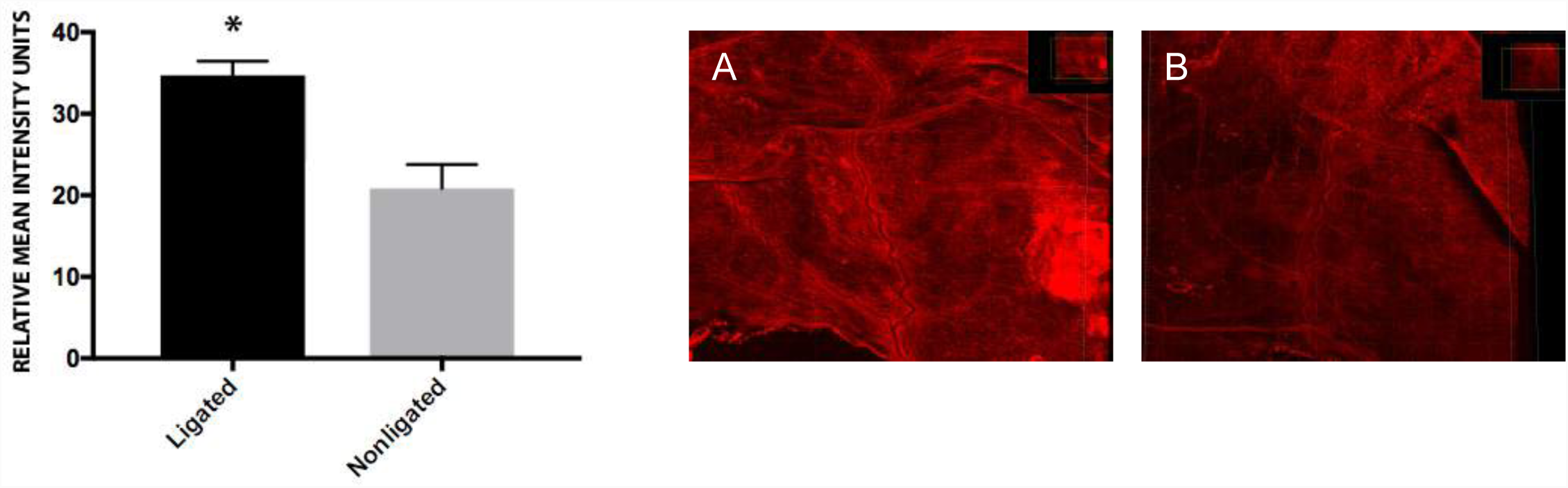
Quantitative effects of cervical lymphatic ligation on lateral meningeal pan-Aβ content. Fig. 4 Legend. Meninges from age-matched ligated and non-ligated TgF344-AD animals were collected and immunostained to Aβ per standard protocol. Full-thickness confocal images were acquired under identical conditions and used to obtain a maximal intensity projection, using standard gain settings. An arbitrary region of 2000×2000voxels was mapped around the base of the middle meningeal artery and analyzed in ImageJ for relative mean intensities across the region. These values provide a quantitative marker of relative Aβ deposition in the meninges between groups. Ligated animals had a statistically significant difference (P=0.0045, Welch’s two-tailed t-test, N=3 in each group) in amyloid content in the region of interest, suggesting that lateral vascular structures around the middle meningeal artery contribute to amyloid efflux, which is impaired in the setting of cervical lymphatic ligation. Representative images of meninges from age-matched ligated (A) and non-ligated (B) animals, stained with 4G8 antibody.

### Pyroglutamate abeta (pE3-Aβ) is avidly taken up by rat meningeal lymphatic vessels

Pyroglutamate Aβ (pE3-Aβ) is a N-terminal truncated and highly neurotoxic form of Aβ. It is more toxic to neuronal and astrocyte cell culture than Aβ42, confers resistance to amyloid degradation, and has direct adverse effects on cell membranes and lipid peroxidation *(31,32).* Soluble oligomers of Aβ are a major feature of cognitive changes mediated at the level of the synapse *(33),* and a primary difference in amyloid composition between normal aging and Alzheimer’s is the prevalence of N-terminal truncated forms *(34).* pE3-Aβ propagates Aβ oligomers in prion-like fashion even when present in very small amounts *(35)* and may be responsible for Aβ seeding across brain regions. It is known to build up in lysosomes of both neurons and glia *(36)* and disrupts hippocampal long-term potentiation *(37).* Though estimates vary depending on quantification methods, prior studies have shown that pE3-Aβ makes up a large fraction of total Aβ in the human Alzheimer’s brain *(38-40),* particularly in pre-amyloid lesions and amyloid plaques where it causes accelerated seed formation for aggregates *(41,42).* Both our data in this study and prior pathology studies *(43)* suggest that pE3-Aβ frequently forms the cores of hippocampal plaques and is found within hippocampal neurons. In a number of new transgenic animal models *(44,45)* intraneuronal pE3-Aβ causes a severe phenotype in the absence of extraneuronal plaques, and thus may act both intracellularly and extracellularly. Given this evidence of its role in Alzheimer’s disease progression, brain clearance pathways associated with pE3-Aβ have major therapeutic implications.

We now report that pE3-Aβ strongly co-localizes with meningeal lymphatic vessels in immunohistochemical preparations of whole meninges, throughout diverse regions of the meninges **[Fig.5]**. The strongest immunohistochemical pE3-Aβ signal was detected around the central venous sinuses and pineal gland, the lateral lymphatic plexus running with the middle meningeal artery (MMA), and the anterior olfactory region. This finding is consistent with our biochemical data which indicate that pE3-Aβ is found in aged TgF344-AD rats at similar high levels in the hippocampus and meninges, and is potentiated by lymphatic blockade.

**Fig. 5.**
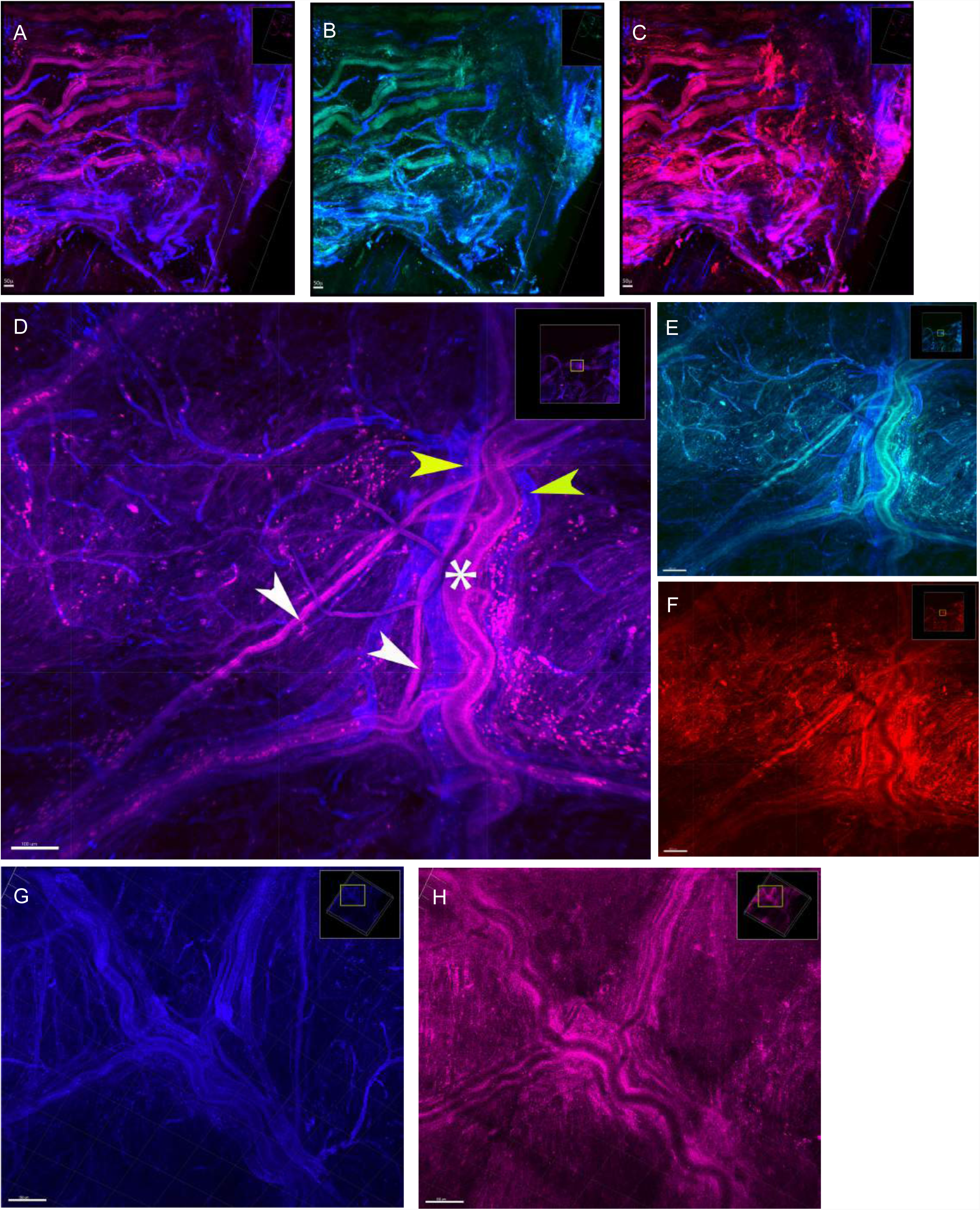
Specific uptake of pE3-Aβ by dural lymphatics and presence of meningeal Mato cells. Fig. 5 Legend. Immunohistochemical labeling to markers of lymphatics co-localize with Aβ and pE3-Aβ. (A) Staining to lymphatic marker Pdpn (purple) and vascular marker CD31 (blue) identifies lymphatics and veins at the border of transverse sinus in TgF344-AD rat following lymphatic ligation. Lymphatics show strong staining to pE3-Aβ (green) and Aβ (red) in (B) and respectively. (D) Close-up of middle meningeal artery and surrounding lymphatic plexus, showing autofluorescent Mato cells. Paired lymphatics (white arrows), veins (yellow arrows), and MMA (asterisk) are shown. (E) Specific lymphatic and periarteriolar staining to pE3-Aβ (green). (F) Strong pan-Aβ signal (red) in lymphatic and periarteriolar distribution. (G) High resolution image of MMA, veins, and lymphatics with punctate perivascular cells around artery and veins. (H) CD68 staining to macrophages using quantum dots shows overlap with punctate cells and perivascular region generally.

The MMA itself (but not meningeal veins) was also strongly positive for pE3-Aβ, raising the possibility of paravascular drainage of pE3-Aβ in the same direction as abutting lymphatic vessels. Interestingly, dura surrounding the MMA had strong pan-Aβ signal, including along the collecting lymphatics and meningeal veins, but the intensity of pan-Aβ staining within the MMA itself was much less, limited to a segmental amyloid pattern which is typical of cerebral amyloid angiopathy (CAA) *(46).* CAA is common in leptomeningeal vessels, but CAA in pachymeninges is less well studied. In most rat specimens, we observed dual lymphatic vessels and dual meningeal veins flanking the MMA, with some variability in morphologies. The lymphatic plexus in this lateral meningeal region was recently reported to be present early in postnatal developmental and extends from the foramen spinosum to the cranial vault *(47,48).* This lymphatic plexus is particularly interesting in light of recent anatomical studies showing that the human MMA is invested with a venous sinus sheath *(49,50).* Thus the presence of lymphatic vessels running with meningeal veins and venous sinuses appears to be a general phenomenon.

An unexpected finding was the presence of large numbers of autofluorescent perivascular cells surrounding the middle meningeal vessels. Because the identity of these cells was unclear, we restained our original preparations using quantum dot tagged immunolabeling and confirmed that they were CD68+ macrophages. These cells were also positive for the lymphatic marker podoplanin, previously reported as a common marker of inflammatory macrophages *(51).* The fact that these macrophages appear autofluorescent on multiple channels suggests that they are “Mato cells” or fluorescent granular perithelial cells *(52).* Perivascular macrophages are associated both with Aβ clearance *(53,54)* as well as lymphangiogensis *(55)* which makes their presence in this location noteworthy.

### Glutamyl cyclase, the enzyme which converts Aβ to pE3-Aβ, co-localizes with lymphatic markers

Glutamyl cyclase (QC) is the enzyme required for conversion of N-terminal truncated Aβ isoforms to pE3-Aβ. QC levels are highly correlated with pE3-Aβ load, which is a more reliable predictor of cognitive decline using MMSE than unmodified Aβ *(56,57).* Of relevance to our study, prior studies have shown that QC gene expression is upregulated 16-fold in lymphatic endothelial cells (LEC) compared to blood endothelial cells (BEC) *(58),* but the relevance of this observation was previously unknown. Although our RNAseq analysis did not detect QC dysregulation among different lymphatic cells *in vitro,* QC has been reported by others to be expressed at higher levels in Alzheimer’s disease (with overall enzyme activity similar to controls). Notably, inhibition of QC has been shown to attenuate pE3-Aβ and classic Alzheimer’s pathology and is a current therapeutic target for Alzheimer’s disease *(59).*

After observing increased relative pE3-Aβ signal in lymphatic vessels, we immunocytochemically confirmed QC expression in isolated lymphatic cells from TgF344-AD rats, and then stained whole meninges for the presence of QC alongside lymphatic markers. We found that QC was expressed in both meningeal lymphatics and arteries, and strongly co-localized with lymphatic markers Lyve1 and Pdpn along the central venous sinuses **[Fig.6]**. This suggests that in addition to neurons, lymphatic vessels are a potential target for QC inhibition. However, it is unknown if the kinetics of pE3-Aβ and Aβ transport are different in lymphatic cells, or if there is any clearance advantage to this Aβ conversion within the lymphatic compartment. If that were proven to be the case, then blocking QC in this compartment might actually impair clearance.

**Fig. 6.**
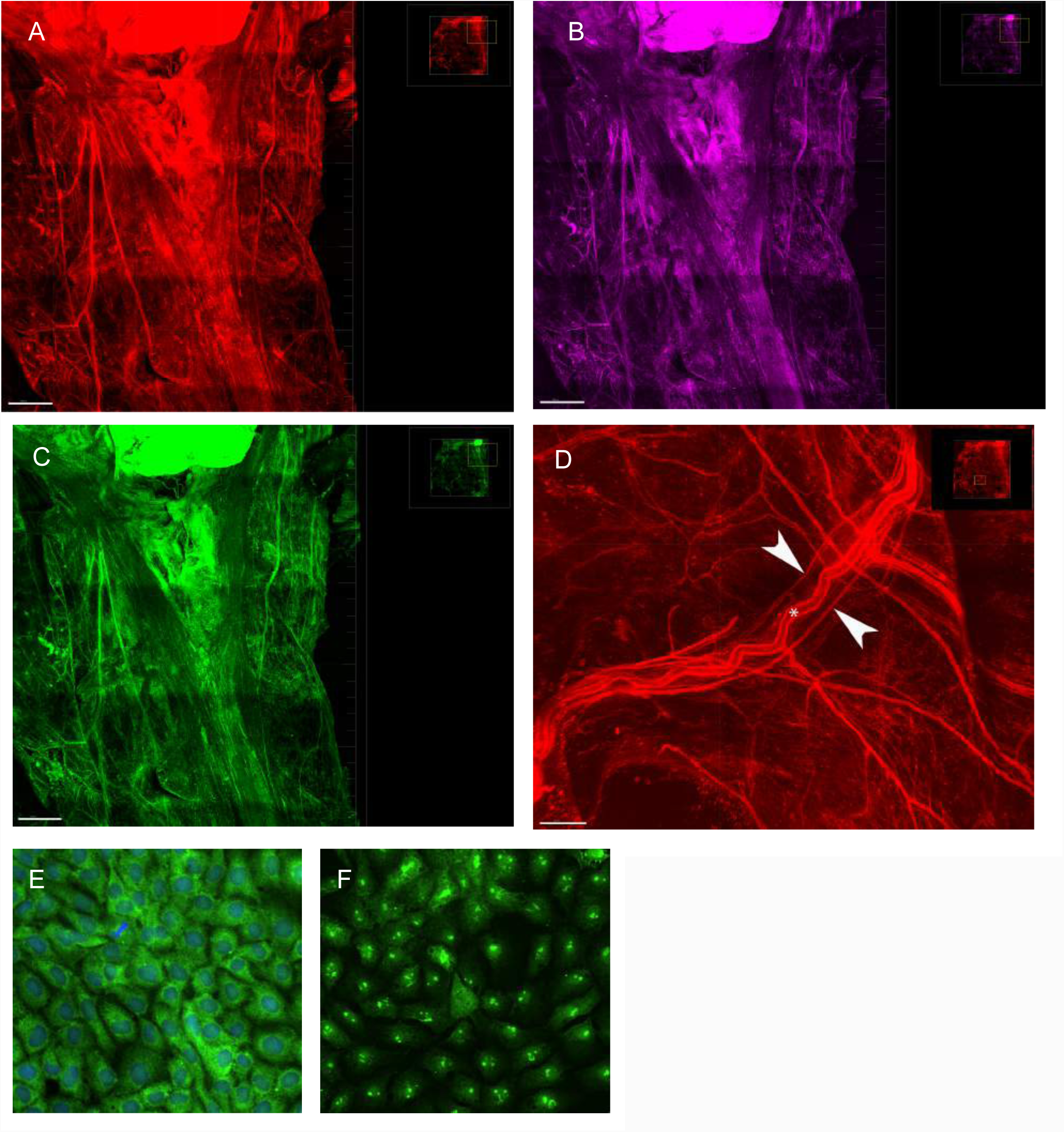
Glutamyl cyclase is enriched in meningeal lymphatic cells and vessels. Fig. 6 Legend. (A) Immunohistochemical labeling to glutamyl cyclase (QC) around the confluence of sinuses and pineal gland, showing co-labeling to lymphatic markers (B) Pdpn and (C) Lyve1. Staining of lateral meninges (D) also shows strong QC staining of the MMA and flanking lymphatic collecting ducts. ICC with isolated rat lymphatic primary cells BrLym1 similarly shows strong signal (E) for QC and (F) lymphatic marker Prox1.

### pE3-Aβ and Aβ accumulate at early stages in meninges, in parallel with increases in brain tissues

In this experiment, our goal was to examine which areas showed the earliest amyloid changes, and also how the effects varied by age and disease status. Consistent with data from our immunohistochemical analysis, biochemical analyses of meninges using sensitive ELISA and electrochemiluminescence assays indicate that the dura and its associated vasculature accumulate pE3-Aβ and Aβ starting in early adulthood [Table 1], starting after the age of 4 months. The blood-CSF barrier (arachnoid), meninges, and olfactory region showed early and relatively large accumulation of both pE3-Aβ and Aβ42, which significantly increased with age. In parallel with changes in meninges, we observed increases of pE3-Aβ and Aβ42 in cognitively important brain regions including the hippocampus. These data suggest that both the pachymeninges (dura) and the leptomeninges are linked with clearance of amyloid that originates elsewhere in the brain. The superficial and deep lymph nodes did not show a significant difference in pE3-Aβ with respect to Alzheimer’s genotype, but did show a significant increase with age; with Aβ42, there were increases both with respect to Alzheimer’s genotype and age, but these were relatively small in magnitude compared to increases elsewhere. The lack of greater amyloid build-up in cervical lymph glands was somewhat unexpected, but one possibility is that amyloid may be actively broken down or exported from this location.

**Table 1A.**
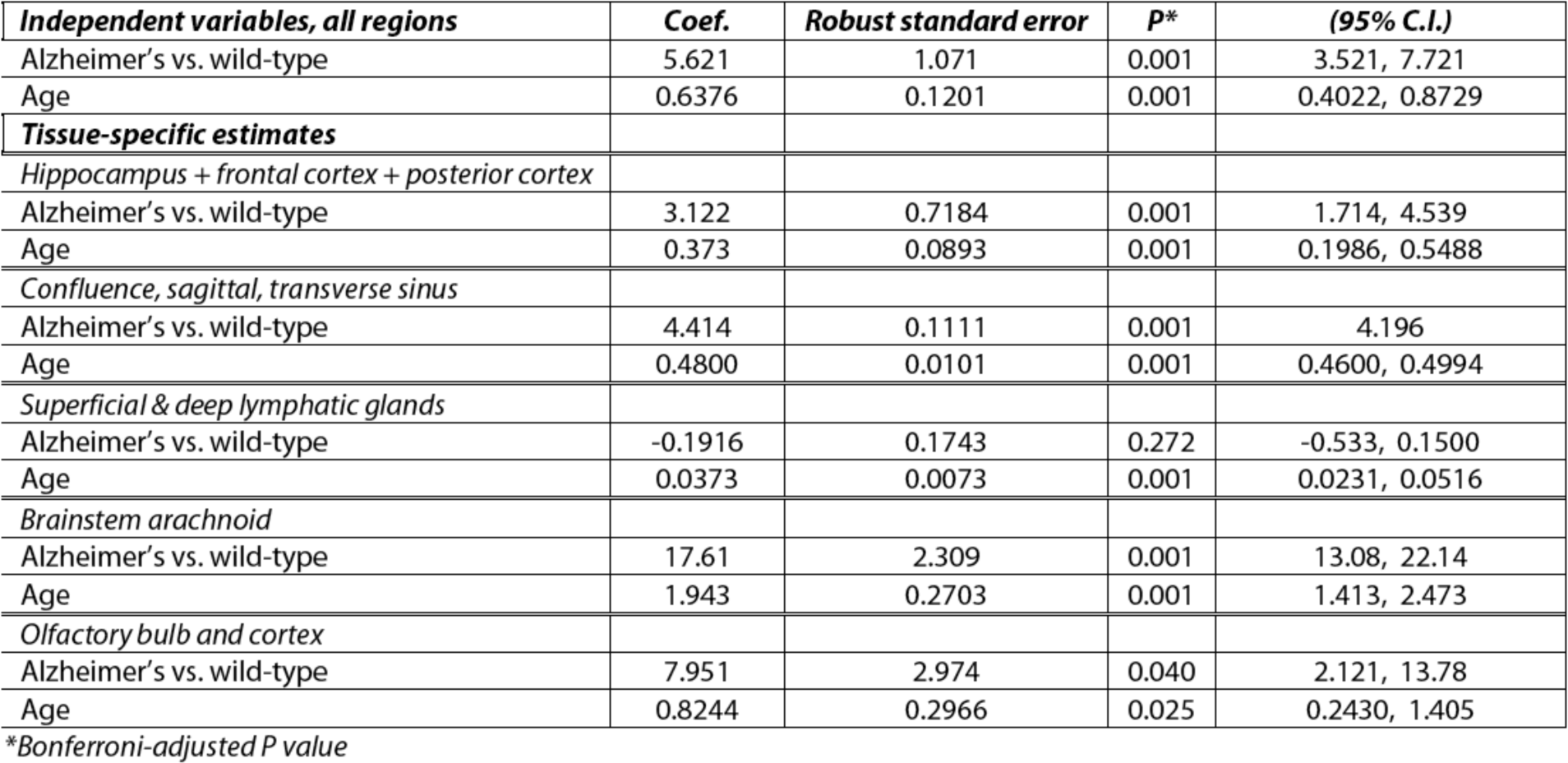
Effects of genotype, age, brain region on pE3-Aβ pathology

| *Independent variables, all regions* | *Coef.* | *Robust standard error* | *P\** | *(95% C.I.)* |
| --- | --- | --- | --- | --- |
| Alzheimer's vs. wild-type | 5.621 | 1.071 | 0.001 | 3.521, 7.721 |
| Age | 0.6376 | 0.1201 | 0.001 | 0.4022, 0.8729 |
| ***Tissue-specific estimates*** |  |  |  |  |
| *Hippocampus + frontal cortex + posterior cortex* |  |  |  |  |
| Alzheimer's vs. wild-type | 3.122 | 0.7184 | 0.001 | 1.714, 4.539 |
| Age | 0.373 | 0.0893 | 0.001 | 0.1986, 0.5488 |
| *Confluence, sagittal, transverse sinus* |  |  |  |  |
| Alzheimer's vs. wild-type | 4.414 | 0.1111 | 0.001 | 4.196 |
| Age | 0.4800 | 0.0101 | 0.001 | 0.4600, 0.4994 |
| *Superficial & deep lymphatic glands* |  |  |  |  |
| Alzheimer's vs. wild-type | -0.1916 | 0.1743 | 0.272 | -0.533, 0.1500 |
| Age | 0.0373 | 0.0073 | 0.001 | 0.0231, 0.0516 |
| *Brainstem arachnoid* |  |  |  |  |
| Alzheimer's vs. wild-type | 17.61 | 2.309 | 0.001 | 13.08, 22.14 |
| Age | 1.943 | 0.2703 | 0.001 | 1.413, 2.473 |
| *Olfactory bulb and cortex* |  |  |  |  |
| Alzheimer's vs. wild-type | 7.951 | 2.974 | 0.040 | 2.121, 13.78 |
| Age | 0.8244 | 0.2966 | 0.025 | 0.2430, 1.405 |

\*Bonferroni-adjusted P value

**Table 1B.** Effects of genotype, age, brain region on Aβ42 pathology

| *Independent variables, all regions* | *Coef.* | *Robust standard error* | *P\** | *(95% C.I.)* |
| --- | --- | --- | --- | --- |
| Alzheimer's vs. wild-type | 1991 | 449.8 | 0.001 | 1109, 2873 |
| Age | 214.3 | 52.30 | 0.005 | 112.8, 317.8 |
| ***Tissue-specific estimates*** |  |  |  |  |
| *Hippocampus + frontal cortex + posterior cortex* |  |  |  |  |
| Alzheimer's vs. wild-type | 2989 | 369.5 | 0.001 | 2264, 3713 |
| Age | 326.5 | 40.98 | 0.001 | 246.1, 406.8 |
| *Confluence, sagittal, transverse sinus* |  |  |  |  |
| Alzheimer's vs. wild-type | 41.92 | 6.703 | 0.001 | 28.78, 55.06 |
| Age | 4.199 | 0.6673 | 0.001 | 2.892, 5.507 |
| *Superficial & deep lymphatic glands* |  |  |  |  |
| Alzheimer's vs. wild-type | 12.61 | 2.044 | 0.001 | 8.605, 16.62 |
| Age | 1.26 | 0.2041 | 0.001 | 0.6850, 1.665 |
| *Brainstem arachnoid* |  |  |  |  |
| Alzheimer's vs. wild-type | 2211 | 448.4 | 0.001 | 1332, 3090 |
| Age | 241.5 | 50.02 | 0.001 | 143.4, 339.5 |
| *Olfactory bulb and cortex* |  |  |  |  |
| Alzheimer's vs. wild-type | 1389 | 37.88 | 0.001 | 1315, 1463 |
| Age | 136.8 | 3.822 | 0.001 | 129.4, 144.3 |

\*Bonferroni-adjusted P value

Using a generalized linear mixed model approach, overall we found statistically significant effects of age, genotype, and region on accumulation of Aβ. Measurements of pE-Aβ and Aβ42 were regressed on disease status and age, and stratified on tissue origin type with clustering to account for within-animal correlation using separate generalized linear models of the form G(E(Y)) = *α* + ^*K*^Σ_k=1_ *β*_*k*_ *X*_*ik*_ where G(E(Y)) is the function of an expected value of Y and Y∼F (Gaussian distribution). G is represented by an identity link function under the distribution of F. Coefficients were calculated according to *a priori* region groupings with residual stratification where needed. Models were fit using a Newton-Raphson (maximum likelihood) optimization E(Y) = *β*_*k*_ *X*_*ik*_, y∼Normal. Averaged cross all regions, Alzheimer’s animals were associated with an increase of 22pg/mg tissue of pE3-Aβ protein compared to wild-type (P = 0.004). Similarly, averaged across all regions, an increase in animal age of one month was associated with pE3-Aβ increase of 1.5pg/mg tissue (P = 0.001).

### Lymphatic ligation accelerates Alzheimer’s pathology in meninges & cognitively important brain regions

Superficial and deep lymphatic chains were microsurgically obstructed bilaterally using titanium microclips **(fig. S4)** for variable periods of time in rats of different ages. The purpose was to determine if cervical ligation would accelerate the expected progression of Alzheimer’s pathology, which was assessed using stereological and biochemical techniques. For our biochemical analysis, all ligated animals were TgF344-AD rather than wild-type, in order to test the specific effect of ligation in animals which inevitably become symptomatic. Several old rats (15 months) did not tolerate this procedure well, exhibiting a phenotype of Foldi’s lymphatic encephalopathy manifest as lethargy and head droop; these animals were humanely euthanized within a week of ligation. Younger rats (5 months) tolerated the procedure for extended periods.

#### Stereological evidence of increased hippocampal plaques

After determining that meninges displayed increased amyloidosis post-ligation, we examined upstream brain regions for evidence of impaired clearance, hypothesized to be a result of a “traffic jam” of Aβ clearance at the meninges. Using unbiased stereological sampling, we systematically compared hippocampi of ligated and non-ligated animals for area fraction of Aβ plaques. IHC was performed on fixed tissue obtained from ½ of the brain. Qualitatively, in both ligated and non-ligated TgF344-AD animals, we found that pE3-Aβ localized along blood vessels in a tram-track pattern starting at 8 months and was also present in a stippled pattern inside hippocampal neurons. This pathology was later accompanied by tau starting at approximately 15 months, as demonstrated with staining to tau/PHF. Quantitative stereological measurements of total plaque areas showed a statistically significant effect between ligated and non-ligated groups **[Fig 7]**.

**Fig. 7.**
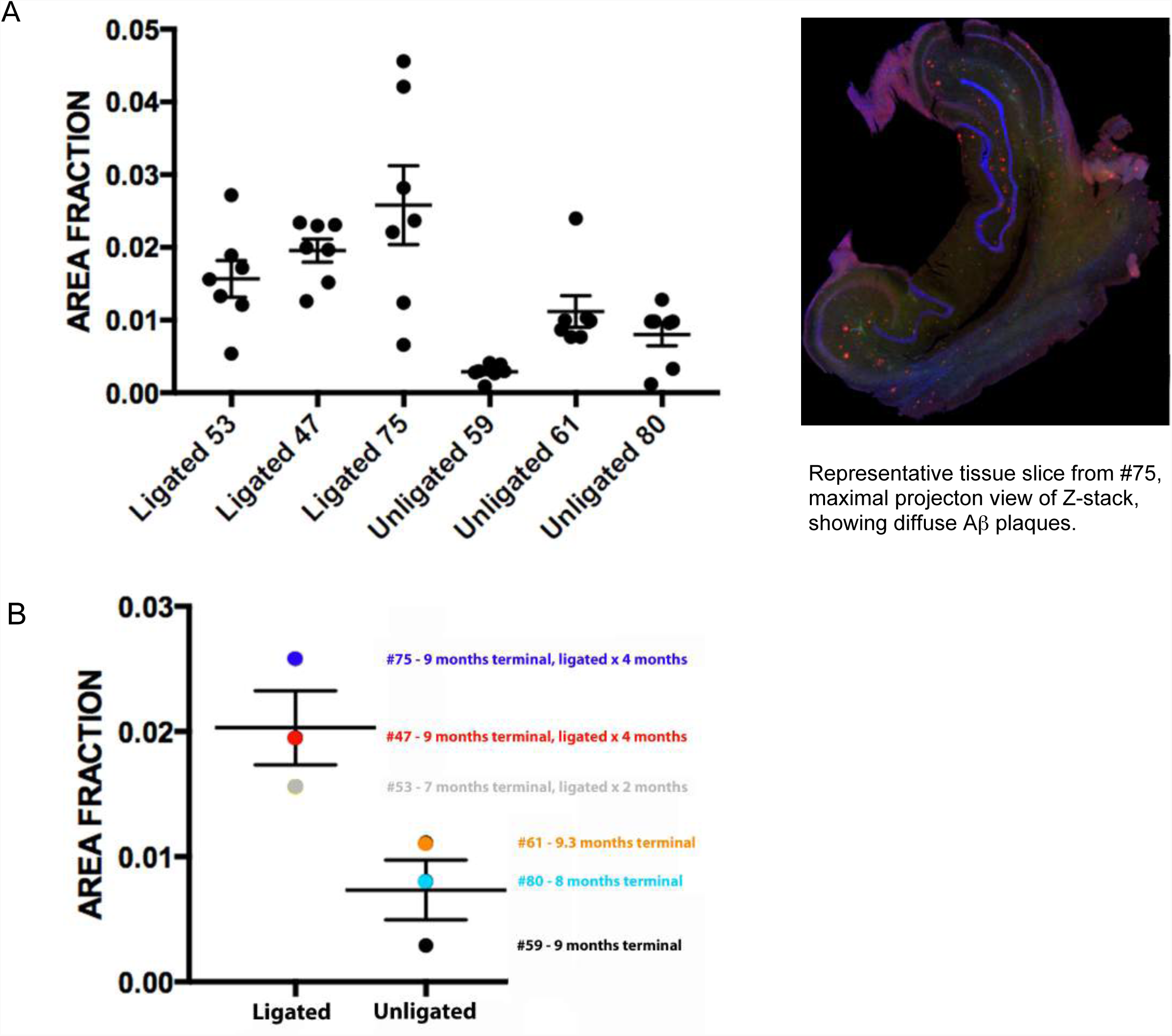
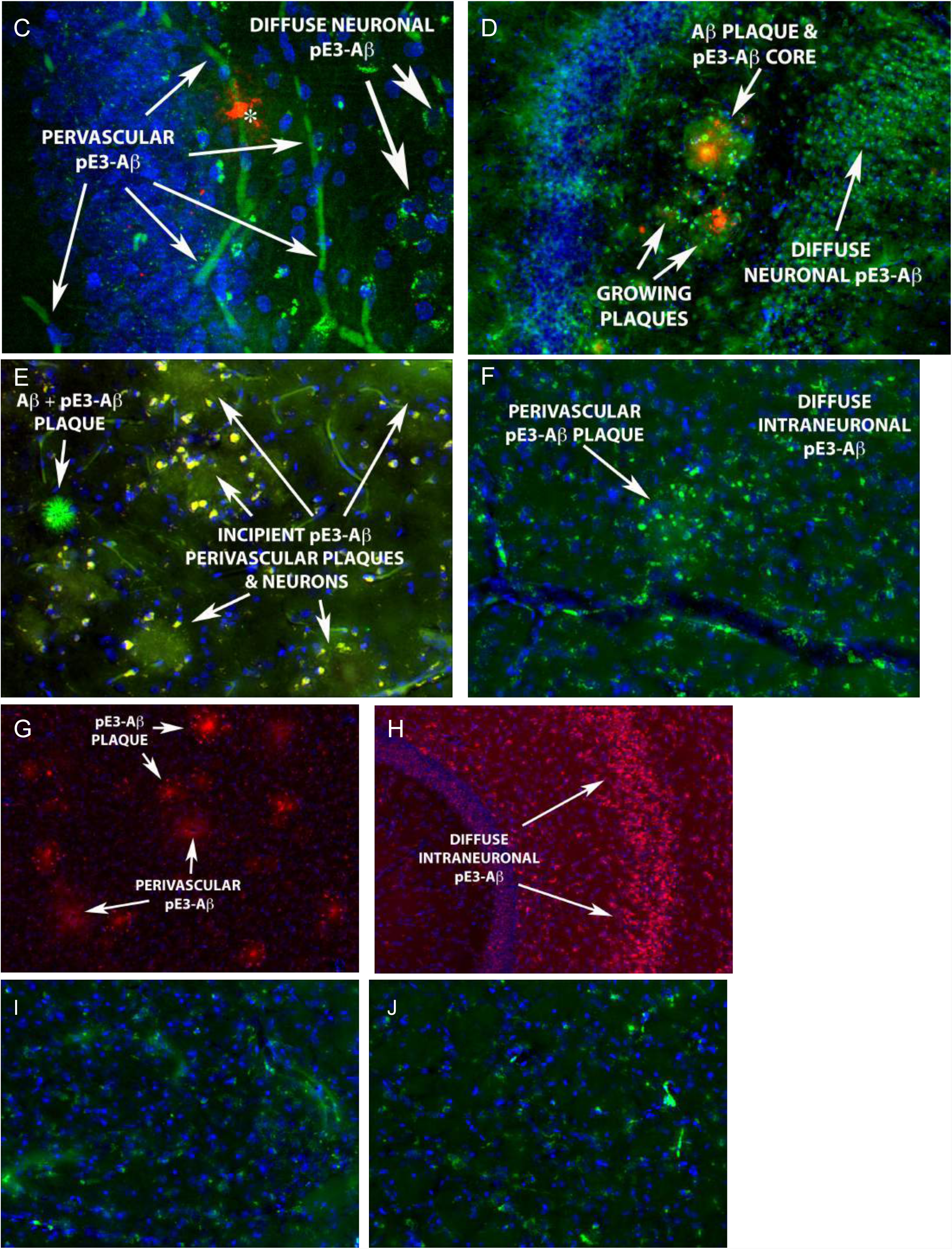
Hippocampal pathology: microvascular, neuronal, and extracellular deposition of pE3-Aβ. Fig. 7 Legend. Comparing area fractions of amyloid plaques in hippocampus using unbiased stereology, we found a significant difference (P=0.004) between ligated and non-ligated groups (two-tailed t test). Mean area fractions by animal with individual tissue counts (A) and by group (B) are shown. Representative images (C,D) of perivascular, intraneuronal, and extracellular pE3-Aβ in hippocampus of ligated animal #47 at 9 months, showing tram-track morphology of capillary amyloid angiopathy; and (E) ligated animal #43 at 18 months. Hippocampus of unligated TgF344-AD rat at 20 months (F) with larger penetrating vessel shows similar perivascular deposits. Unligated TgF344-AD rat at 8 months (G) shows incipient perivascular pE3-Aβ, which by 23 months is present throughout the hippocampal neurons. Staining to tau paired helical filaments (H) and phosphorylatyed tau (I) is evident starting at 12-15 months, shown here in a 20 month unligated animal.

#### Quantitative biochemical assays to Aβ and pE3-Aβ

To augment our IHC analysis, we performed biochemical assays on fresh tissue extracts from the other ½ of each brain, using ELISA and MSD-ECL. Results show a significant effect of both ligation and age on brain levels of pE3-Aβ and Aβ in hippocampus and cortex. This quantitative difference in pE3-Aβ and Aβ content implies that cervical lymphatic ligation potentiates intracranial disease in multiple brain regions important for cognition. The effect on pE3-Aβ was most significant, followed by Aβ42. However, effects of ligation on Aβ40 were not significant, which suggests that other clearance mechanisms may compensate. It has been shown in prior studies that Aβ40 is preferentially cleared by the blood-brain barrier, and this differential clearance may explain the observed results.

GLM analysis **[Table 2]** shows that cervical lymphatic ligation resulted in a statistically significant increase in pE3-Aβ across multiple brain regions (hippocampus, frontal cortex, posterior cortex). The multivariable GLM is adjusted for age at sacrifice and length of ligation (correlation on animal and region, family=Gaussian, link=log). The frontal region includes motor and olfactory areas, and posterior cortex includes parietal and occipital areas. Levels of pE3-Aβ by region represent total pE3-Aβ combined from the Triton and guanidine fractions. P-values are Bonferroni-corrected for multiple comparisons.

**Table 2A.**
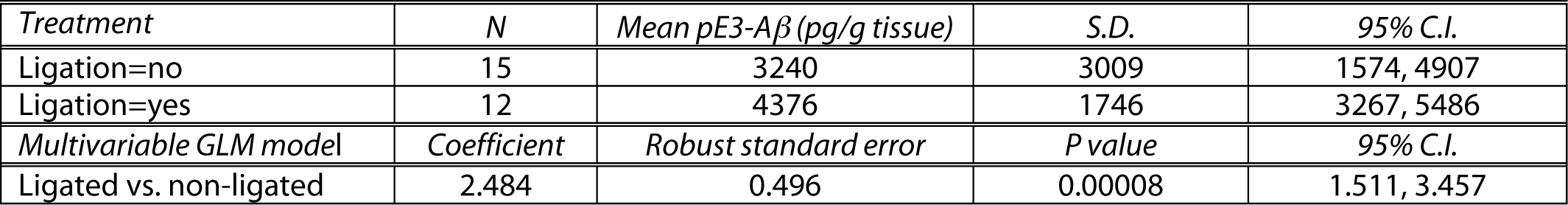
Comparison of pE3-Aβ following lymphatic ligation (Hippocampus + Frontal Cortex + Posterior Cortex)

**Table 2B.** Mean pE3-Aβ by anatomical region for ligated & non-ligated animals

| Anatomical region | N | Mean pE3-Aβ (pg/g tissue) | S.D. | 95% C.I. |
| --- | --- | --- | --- | --- |
| Hippocampus | 9 | 4687 | 3034 | 2354, 7019 |
| Frontal cortex | 9 | 3011 | 1765 | 1654, 4367 |
| Posterior cortex | 9 | 3539 | 2672 | 1484, 5593 |
| Arachnoid | 9 | 19875 | 14040 | 9083, 30667 |
| Choroid plexus | 9 | 40442 | 23889 | 22079, 58806 |
| Repeated measures ANOVA | Partial SS | Df | F | P |
| Any region | 9.452 x 10<sup>9</sup> | 4 | 16.64 | 0.0000006 |
| Pairwise comparisons (post hoc) | Uncorrected P value |  | Mean difference | 95% C.I. |
| Choroid plexus vs. Hippocampus | 0.0020 |  | 35756 | 17345, 54166 |
| Arachnoid vs. Hippocampus | 0.0117 |  | 15188 | 4309, 26067 |
| Frontal cortex vs. Hippocampus | 0.1759 (ns) |  | 1676 | -855, 4207 |
| Posterior cortex vs. Hippocampus | 0.4070 (ns) |  | 1148 | -1713, 4009 |

**Table 2C.** Comparison of Aβ42 following lymphatic ligation (Hippocampus + Frontal Cortex + Posterior Cortex)

| Treatment | N | Mean Aβ42 (pg/g tissue) | SD | 95% C.I. |
| --- | --- | --- | --- | --- |
| Ligation=no | 15 | 2595411 | 2601219 | 1154903, 4035918 |
| Ligation=yes | 12 | 3979192 | 3937768 | 1477253, 6481130 |
| Multivariable GLM model | Coefficient | Robust standard error | P value | 95% C.I. |
| Ligated vs. non-ligated | 1.983 | 0.353 | 0.00002 | 1.291, 2.675 |

**Table 2D.** Comparison of Aβ40,42 following lymphatic ligation (Hippocampus + Frontal Cortex + Posterior Cortex)

| Treatment | N | Mean Aβ40,42 (pg/g tissue) | SD | 95% C.I. |
| --- | --- | --- | --- | --- |
| Ligation=no | 30 | 1951839 | 2216713 | 1124105, 2779573 |
| Ligation=yes | 24 | 2741865 | 3146701 | 1413130, 4070600 |
| Multivariable GLM model | Coefficient | Robust standard error | P value | 95% C.I. |
| Ligated vs. non-ligated | 1.313 | 0.346 | 0.0009 | 0.636, 1.991 |
| Anatomical region | N | Mean Aβ40,42 (pg/g tissue) | SD | 95% C.I. |
| Hippocampus | 18 | 1188863 | 2071142 | 158908, 2218818 |
| Frontal cortex | 18 | 642005 | 1061047 | 114359, 1169652 |
| Posterior cortex | 18 | 2875249 | 2705893 | 1529639, 4220858 |
| Arachnoid | 18 | 1306927 | 1558893 | 531707, 2082146 |
| Choroid plexus | 18 | 2726709 | 3305906 | 1082721, 4370697 |
| Repeated measures ANOVA | Partial SS | Df | F | P value |
| Any region | 7.126 x 10<sup>13</sup> | 4 | 3.53 | 0.154 |

In these experiments, our goal was to examine the effects of ligation, as well as age and length of ligation, with respect to amyloid deposition. Notably, mean levels of pE3-Aβ increased post-ligation in all brain regions tested and also increased with age. This post-ligation effect suggests that the meningeal lymphatic system is necessary for brain clearance of pE3-Aβ, and its absence is not fully compensated by the blood-brain barrier or blood-CSF barrier. Both lymphatic ligation and aging were highly significant effects. Independent of age, ligation results in a mean 5.3-fold increase in total tissue fraction of Aβ (95% C.I. 3.0-9.3, P=0.000002). For age, given a one month increase in age at the time of tissue collection, the total tissue fraction increases by 14% (95% CI 10-18%, P=0.000003) or RR=1.14, 95% CI 1.10 to 1.18.

After determining that cervical lymphatic ligation was highly correlated with pE3-Aβ increases in the brain, we wanted to determine if there were regional differences in pE3-Aβ accumulation across this observed age range (7-19 months) in TgF344-AD animals, whether or not they were ligated. Pooling data from ligated and non-ligated animals, the highest mean pE3-Aβ levels were found in arachnoid and choroid plexus, which were significantly enriched in pE3-Aβ but had comparable amounts of Aβ40/42 compared to hippocampus and cortical regions which contain capillaries of the blood-brain barrier. This is important, because it suggests that lymphatics work synergetically with the CSF-blood barrier to clear pE3-Aβ.

Similar to results with pE3-Aβ, using the GLM approach we found that Aβ42 levels in brain also significantly increased post-ligation. Aβ42 peptide measured by quantitative immunodetection was over 800x more concentrated overall than pE3-Aβ levels. As with pE3-Aβ, mean levels of Aβ42 significantly increased post-ligation in all brain regions. Unlike results with pE3-Aβ, however, we did not detect a statistically significant difference in mean Aβ42 levels by subregion (P=0.59), despite the significant effect of ligation overall, suggesting that Aβ42 is more evenly dispersed throughout the brain.

In the case of Aβ40, we did not detect a statistically significant effect of lymphatic ligation. Given that Aβ40 is preferentially transported across endothelial cells at the blood-brain barrier, this result was not entirely unexpected; Aβ40 transport at the blood-brain barrier may compensate for loss of transport attributable to blocked lymphatic ligation. Since we showed *in vitro* that lymphatics transport Aβ40, an alternative efflux route is the most plausible explanation. Similarly, we did not detect significant region-specific variation in Aβ40 (p=0.11) in regions surveyed. Interestingly, when Aβ40 and Aβ42 were considered altogether, the effect of ligation on amyloid burden remained statistically significant, suggesting that Aβ42 dominates.

### FocusDeep panoramic images of whole-mount human meninges reveal dural lymphatic networks

Given our compelling findings with whole-mount rat meninges, we extended this approach to analysis of the structure and function of human meninges. However, imaging the human meninges presents a number of unique challenges. Among these are the opacity, large surface area, and thickness of this tissue (which ranges from 1-3mm in lateral regions up to 10-20mm at the falx and central sinuses), the diverse composition of dense connective tissue that attracts non-specific antibody binding (including collagen, elastin, laminin, and glycosaminoglycans), and the high prevalence of autofluorescent deposits with aging. To simplify matters, we started by preparing whole-mounts of specimens from human neonatal dura, which are smaller and thinner than adult dura. Our first objective was determining the topology of dural lymphatics, which was achieved using post-fixation clearing techniques with fast mosaic confocal scanning.

A variety of clearing techniques have been developed for brain, but few of these are appropriate for human meninges. In general, tissue clearing works by one of two physical principles: removal of opaque lipid and protein components (by solvation, detergents, or electrophoresis), or refractive index matching *(60).* Clearing techniques with organic solvents render the tissue fragile and are often not compatible with immunofluorescent labeling. Thus we considered a number of aqueous methods based on refractive index matching and hyperhydration including SeeDB *(61),* ScaleS *(62),* FRUIT *(63),* and UbasM *(64).* The SeeDB method had been used successfully for connective tissues *(65)* and was our initial choice. However, FocusClear is a similar refractive index matching agent which we had already used for rat meninges. This is a proprietary mix of diatrizoic acid, DMSO, and other components which hyperhydrate and create a uniform refractive index throughout the tissue *(66).* When pretreated with mild alcohol solvation and mild detergent or bleaching, it is known as FocusDeep.

Using FocusDeep clearing, we obtained the first panoramic images of lymphatic vessels in the human neonatal meninges, using tissue obtained under an IRB-approved protocol. Vessels with classic lymphatic “honeycomb” morphology **[Fig. 8]** stained to Pdpn and Lyve1, markers of lymphatic endothelium, in tissue bordering the sagittal and transverse sinuses. Additional images obtained from the same region had a more venous morphology, which nevertheless stained to both CD31 and Pdpn; the latter could represent either non-specific staining or intermediate stages of lymphatic development, since in mice the meningeal lymphatics develop in the early postnatal period from venous structures *(48).* In humans, nothing is currently known about the timing of meningeal lymphatic development, though these data suggest that at least some some lymphatic vessels are present by the first day of life.

**Fig. 8.**
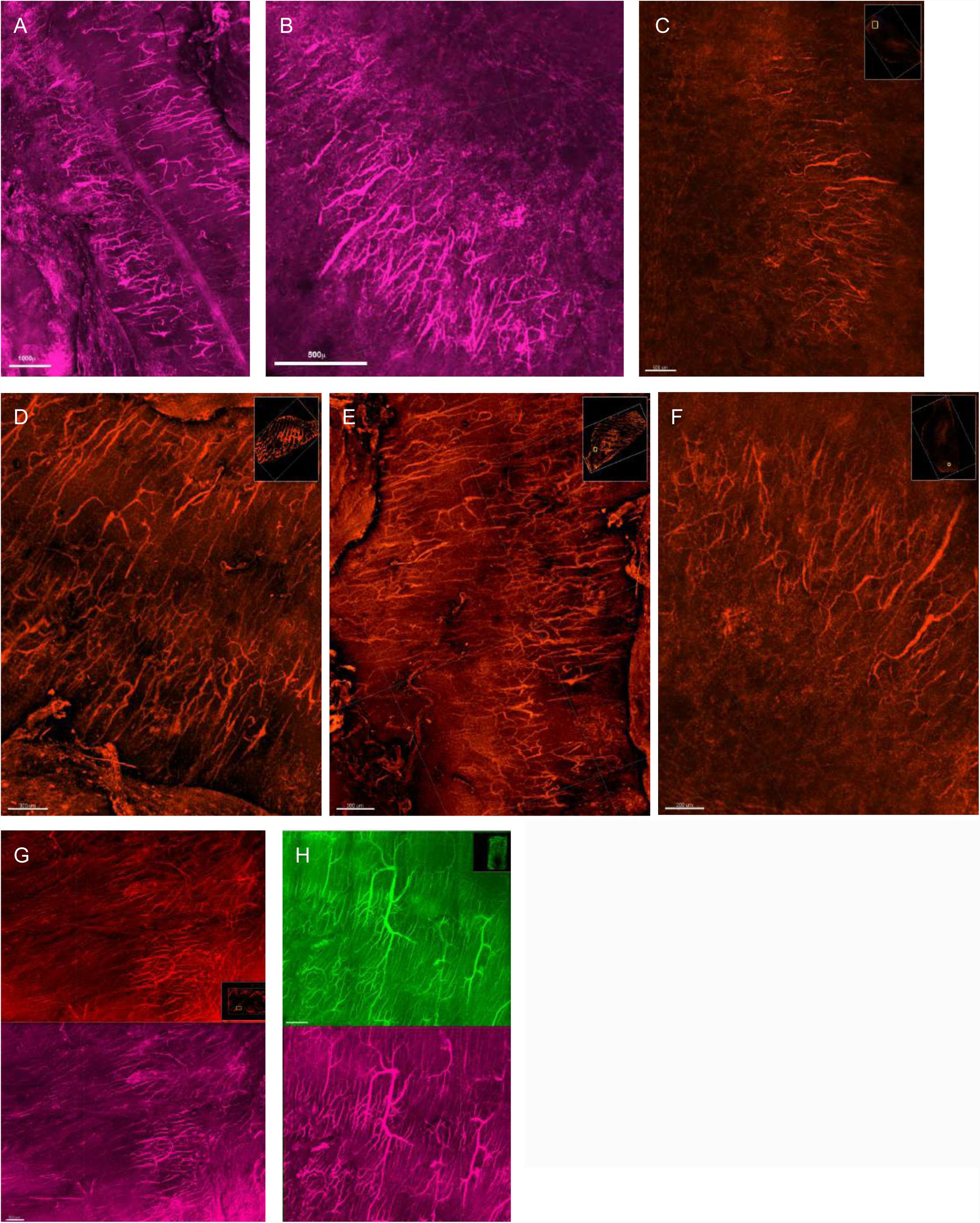
Panoramic images of human dural lymphatic networks. Fig. 8 Legend. The first panoramic images of the human dural lymphatic system, neonatal tissue specimen. Staining to Pdpn (A,B) and Lyve1 (C) along confluence of sinuses. Typical honeycomb morphology with irregular lumen diameter, with Lyve1 staining along sagittal sinus (D,E,F). Venous morphology with multiple branching points in same region (G) with staining to Lyve1 (top) and Pdpn (bottom). Similar venous branching pattern (H) with staining to CD31 (top) and Pdpn (bottom).

### Human Alzheimer’s disease meninges (like rat) show specific uptake of pE3-Aβ in dural lymphatics

Having shown that lymphatic networks are present in human neonatal tissue, we examined human meninges (both sectioned and whole-mount) from Alzheimer’s patients which were treated with FocusClear or FocusDeep, in order to determine if pE3-Aβ or Aβ was present in these vessels. Without clearing, sectioned tissues had an unacceptably high level of background fluorescence, also recently reported by others *(67).* However, cleared tissues showed a much lower level of background autofluorescence from collagen and lipofuscin. Meninges of elderly people are often thin and fragile compared to young adults, due in part to loss of elastin with aging, and the whole-mount meninges we collected along the sagittal sinus were <2mm in thickness and easily mounted on slides with custom silicone gaskets.

We initially observed a variety of CD31+ vessels which were negative for Pdpn or Prox1 and positive for Aβ, confirming prior studies which have reported CAA in the pachymeninges. In addition, we were able to identify a minority of vessels which were positive for multiple lymphatic markers. Interestingly, these vessels were also positive for pE3-Aβ, while venous or arterial structures were positive for Aβ but generally negative for pE3-Aβ **[Fig 9]**. These results support the validity of our data using the TgF344-AD rat and suggest that pE3-Aβ is specifically transported in human dural lymphatics.

**Fig. 9.**
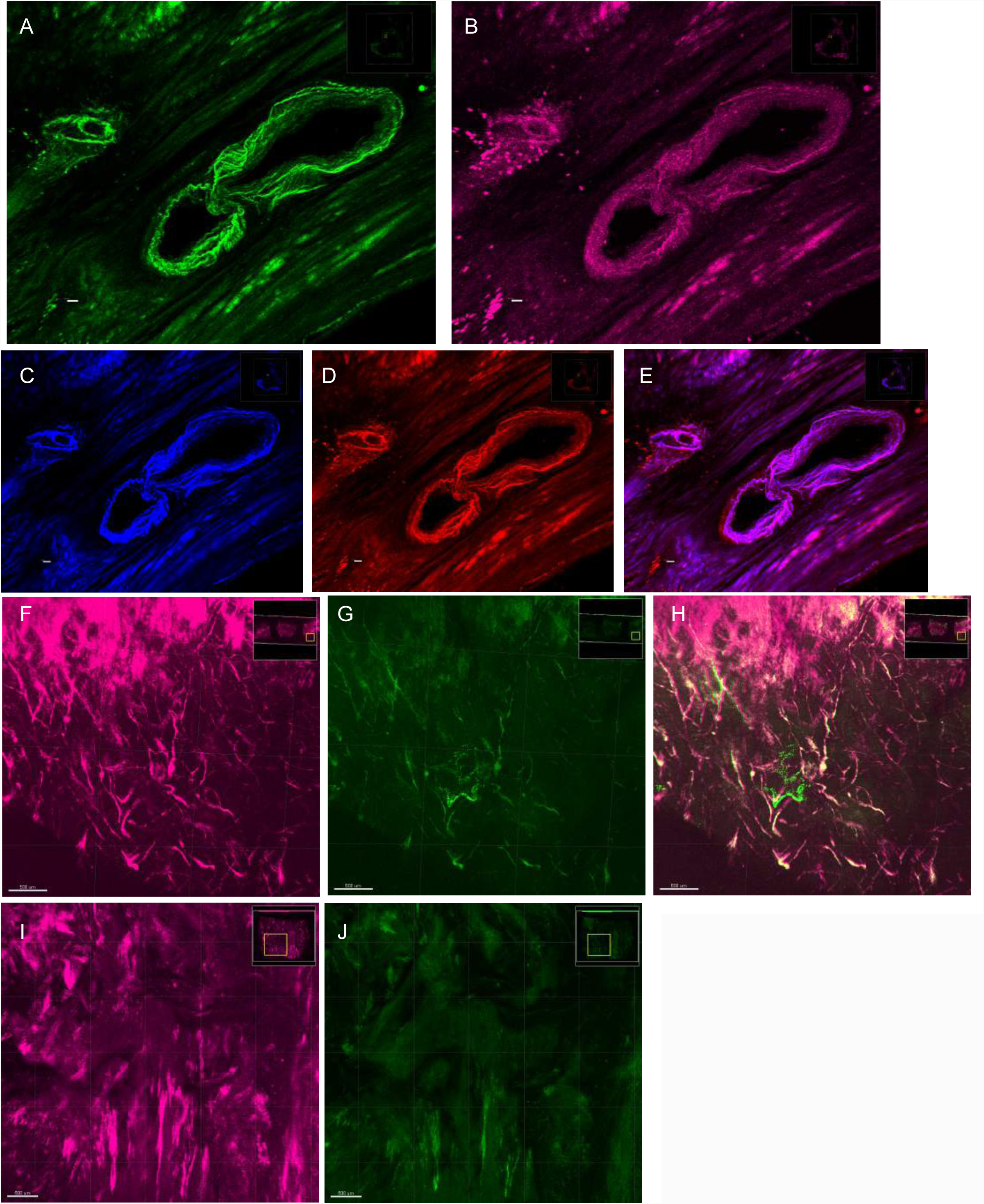
Human dural lymphatics retain pE3-Aβ in Alzheimer’s disease, similar to the TgF344-AD rat. Fig. 9 Legend. Staining to CD31 (A) and Pdpn (B) in FocusClear treated sections shows lumen of lymphatic vessel with surrounding Pdpn+ macrophages as observed in rat tissue, and residual collagen autofluorescence. FocusDeep cleared whole mounts demonstrate Aβ (C), pE3-Aβ (D), and merge (E). Lymphatic morphology shown in staining to Pdpn (F), pE3-Aβ (G), and merge (H). Additional images show Pdpn signal (I) with pE3-Aβ lumenal structures (J).

### Transport of Aβ by lymphatics is energy dependent, and aged Alzheimer’s derived cells transport Aβ less well than young or wild-type cells

Given that energy-dependent processes have been implicated in endothelial transport, we wanted to test the hypothesis that they are also linked with dural lymphatics. The idea of vesicle-mediated endocytosis and transcytosis in endothelial cells was originally pioneered by Nobel laureate George Pallade *(68, 69),* but its physiological role has been generally minimized in favor of paracellular routes. This is even more so with lymphatic endothelia; it is currently accepted dogma in textbooks that lymphatic paracellular transport through discontinuous junctions is the primary means of lymphatic transport of protein and fluid. However, some recent studies have shown that the transcellular route is a major mechanism of solute transport by lymphatic endothelial cells *(70,71).* In support of this mechanism, we also noted in our transmission electron microscopy (TEM) studies (see below) the presence of numerous caveolae and occasional larger vesicles that may be clathrin coated pits. These endocytotic mechanisms are energy dependent through a GTPase called dynamin *(72,73).* Interestingly, one of the seminal early papers in the field of meningeal lymphatics was an elegant TEM study *(74)* which specifically noted frequent dural lymphatic vesicles with coated pits, consistent with clathrin-dependent endocytosis.

Central to our hypothesis of a necessary role for lymphatic clearance in Alzheimer’s disease pathogenesis is the ability of isolated lymphatic cells to transport Aβ. For mechanistic studies to investigate how this putative Aβ transport occurs, we developed a cell sorting protocol to isolate and grow purified lymphatic cells from different groups of animals – young and old, wild-type and Alzheimer’s. Transport from abluminal to luminal side of the membrane was performed using the Transwell model of endothelial transport. In this system, cells are grown on a permeable support interfacing fluid on both sides, simulating the *in vivo* endothelial environment. To demonstrate that lymphatic Aβ transport was energy-dependent, and not solely mediated by paracellular transport, we performed pharmacologic blockade of ATP-dependent processes with 2,4-DNP and specific blockade of <u>A</u>TP-<u>B</u>inding <u>C</u>assette (ABC) transporters, which are implicated as a clearance mechanism of Aβ in transgenic mouse models *(75,76).* We used FITC-conjugated Aβ40/42 rather than pE3-Aβ for *in vitro* transport studies, because our lab already had reliable protocols for preparation of monomeric Aβ40/42 *(77).* In addition to relative ease of preparation, Aβ40/42 is less cytotoxic than pE3-Aβ, permitting a continuous confluent layer of cells with high transendothelial resistance.

Prior to applying Aβ, we verified that lymphatic cell monolayers were continuous by observing with Nomarksi optics, measuring transendothelial resistance (TEER), and obtaining cross-sectional TEM on Transwells which showed the presence of tight junctions. FITC-labeled Aβ40/42 was added to the apical or basal chambers and clearance capacity was measured over time with fluorescence as the read-out using spectrophotometry. Aβ transport was modeled using log regression, which showed statistically significant differences between cells of different age and genotype. We found that cells derived from young wild-type animals transported Aβ significantly better than old cells, and Alzheimer’s cells performed the worst. Moreover, we confirmed that Aβ transport in lymphatics was energy dependent **[Fig. 10]**. The compound Pitstop, which inhibits clathrin-mediated endocytosis, partially blocked transport, and mitochondrial uncoupling agent 2,4-DNP severely impaired transport. The latter effect is expected to be more severe, as it affects both clathrin and caveolin dependent endocytosis and ABC transporter function. These results suggest that functional differences exist between young and old lymphatic cells, and between wild-type and Alzheimer’s cells, and draws a parallel between transport at the blood-brain barrier and meninges.

**Fig. 10.**
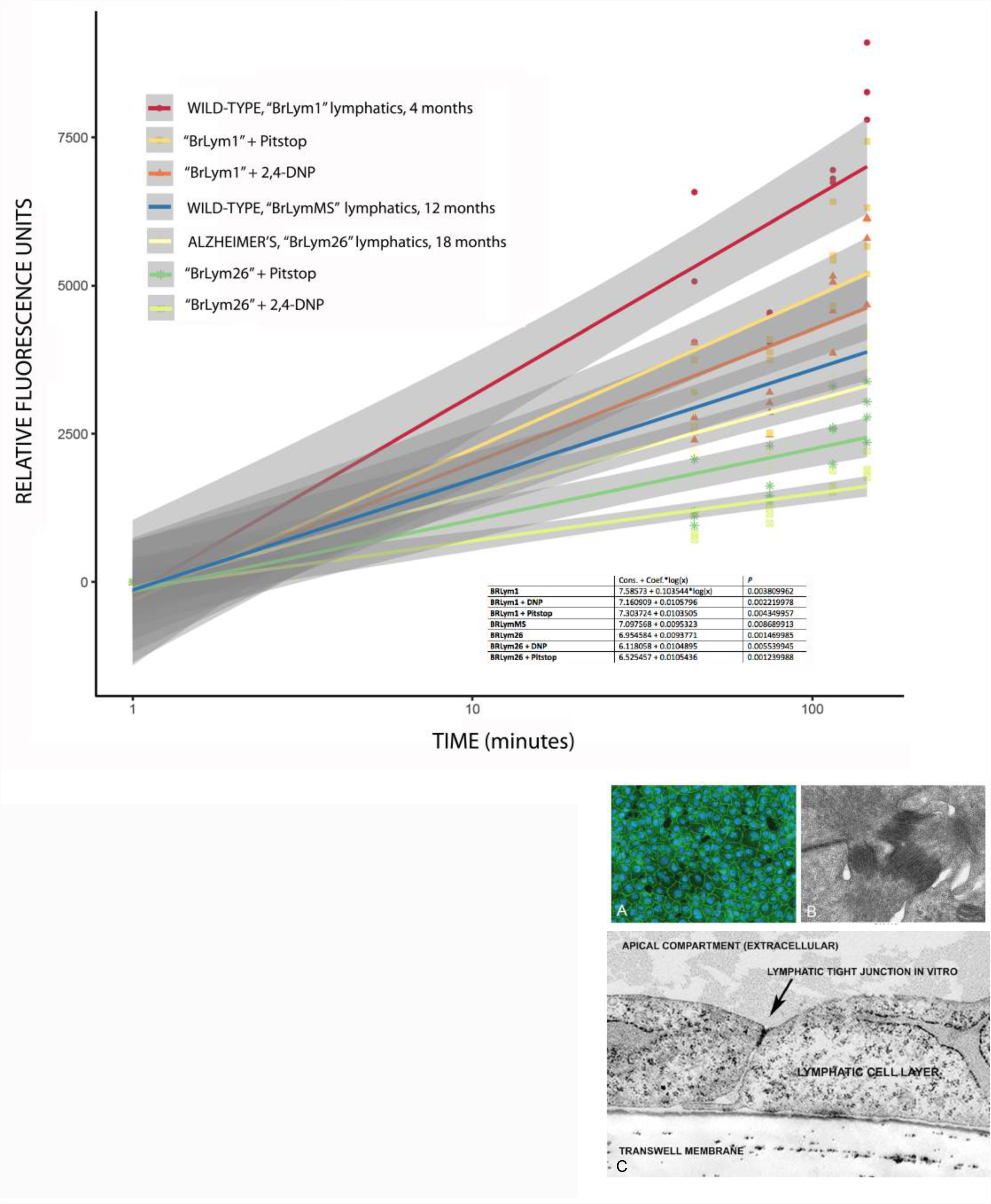
Lymphatic monolayers transport Aβ in an age, disease, & energy dependent fashion. Fig. 10 Legend. Transport of FITC-Aβ40 across lymphatic monolayers in Transwell system. Cells were grown to confluence under standard conditions and 2μM solution was applied. We stained monolayers to junctional marker ZO1 (A) and verified desmosomes and tight junctions with TEM (B,C). TEER values were also used to confirm confluence of live monolayers. There was a statistically significant difference in transport between young and old cells, as well as between wild-type and Alzheimer’s cells. Moreover, inhibition of clathrin-mediated endocytosis with Pitstop partially blocked transport, with a more severe inhibition when 2,4-DNP was used. The latter blocks both clathrin and caveolin dependent endocytosis and ABC transporter mediated endocytosis. The residual transport is presumed to be paracellular or macropinocytosis.

### Energy-dependent <u>A</u>TPase <u>B</u>inding <u>C</u>assette (ABC) transporters are found in meningeal lymphatics

Given our results with Transwell transport assays, which showed that endocytosis is a significant transport mechanism in meningeal lymphatics, we also wanted to verify that functional ABC transporters are operative in lymphatic cells as another component of energy-dependent transport. Prior work had already established that multiple ABC transporters are implicated in Aβ transport at the blood-brain barrier and CSF-blood barrier *(78-81),* but multi-drug transporters have never been reported in meningeal lymphatic cells. However, one prior study *(57)* had reported at the gene expression level that multiple ABC transporters are upregulated in lymphatic endothelial cells (LEC) compared to blood endothelial cells (BEC). In particular, ABCB1 was 50-fold upregulated and ABCC3 was 12-fold upregulated. Our own expression studies (see below) also indicated that multiple types of ABC transporters are highly expressed in dural lymphatic cells including ABCB1, ABCC1, and ABCG2. After immunohistochemically verifying the presence of these ABC transporters in our cell lines, we tested the function of ABC transporters using a flow cytometry assay.

Relative activity of ABC transporters in live meningeal lymphatic cells was compared using age-matched wild-type and Alzheimer’s primary cells “BrLym25” and “BrLym26.” The results of this assay indicate that ABCB1, ABCC1, and ABCG2 are all functionally impaired in Alzheimer’s derived cells compared to age-matched wild-type cells under identical conditions, which suggests that energy-dependent transport at the level of ABC transport may represent another distinct mechanism of clearance in meningeal lymphatics, in addition to energy-dependent endocytosis. Thus, lymphatic endothelia are a potential anatomical target for drugs which modulate multi-drug transporters **[Fig. 11]**.

**Fig. 11.**
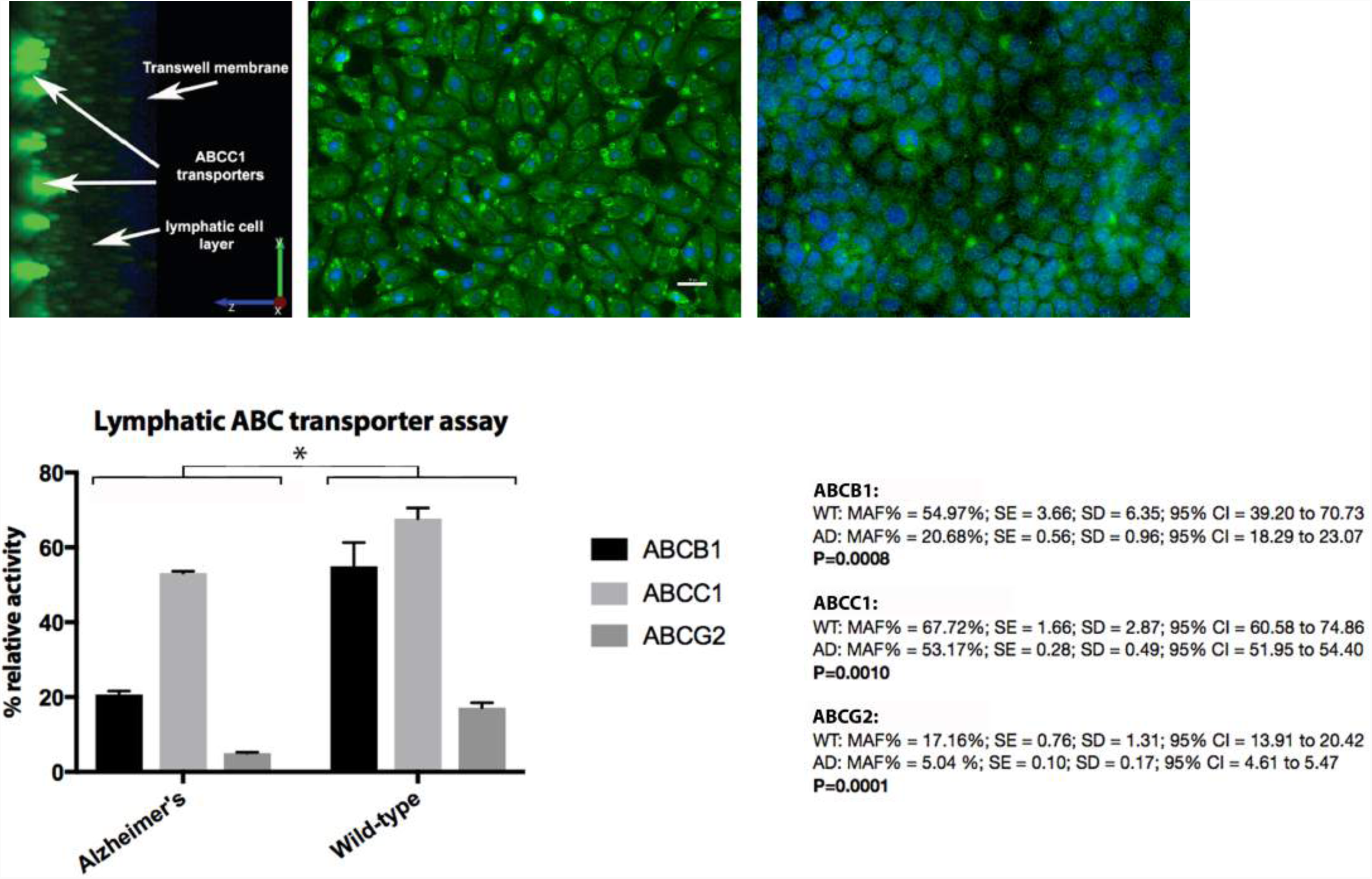
Multiple ATP Cassette-Binding (ABC) multi-drug transporters are active in dural lymphatics. Fig. 11 Legend. ABC activity assay in lymphatic cells, Alzheimer’s vs. wild-type derived. After verifying gene expression and IHC evidence of multiple ABC transporters in lymphatics, including abluminal polarity of expression on Transwells, the age-matched meningeal lymphatic cells were subjected to ABC activity assay using flow cytometry as described. There were significant differences in activity for all ABC transporters tested, consistent with a physiological role in lymphatics. The highest overall activity was in ABCC1, which has been shown in mouse models to be the major Aβ transporter.

### Subcellular pathology in Alzheimer’s derived lymphatic cells is typified by abnormal mitochondria, inclusion bodies, and autophagy

Ultrastructural features of normal dural lymphatic cells include numerous mitochondria, abundant ribosomes and RER, and “button-like” contacts between cells with interspersed open regions which may represent sites of fluid and solute entry of initial lymphatics *in vivo.* We observed flap-like appendages in between adherent cells which may be rudimentary “oak-leaf” interdigitations *in vivo (82).* Wild-type cells imaged with TEM have well-apposed membranes with tight and adherens junctions, and normal appearing organelles. However, age-matched Alzheimer’s genotype cells have less contiguous membranes, bizarre cytoplasmic inclusions, sparse and misshapen mitochondria, and frequent autophagy and mitophagy suggesting a pathological state **(fig. S5)**.

### Evidence for dynamin-dependent transcytosis and paracellular transport in lymphatics

Though plasma membrane vesicles were observed ultrastructurally in lymphatic endothelium for decades, the primary mechanism of lymphatic transmembrane fluid transport has been assumed to be a result of membrane pores and Starling forces. A newer paradigm is that both paracellular and transcellular transport of fluid and protein occurs in lymphatics. To assess the possible contribution of caveolin or clathrin dependent endocytosis as a mediator of Aβ transport, we stained for the presence of clathrin and caveolin in cells isolated from healthy age-matched cells, which showed strong expression to both markers **[Fig. 12]**. In addition, using TEM we ultrastructurally confirmed the prevalence of caveolae and clathrin mediated endocytosis in our cells, which appeared to be particularly concentrated in between the “button-like” intercellular junctions. Extending this observation, we also confirmed *in vivo* that endocytosis occurs in meningeal lymphatics, which appeared particularly robust in wild-type cells. We observed intracellular lymphatic inclusions in meningeal lymphatics from TgF344-AD rats, which were similar in appearance to what we had observed *in vitro* in our cell lines.

**Fig. 12.**
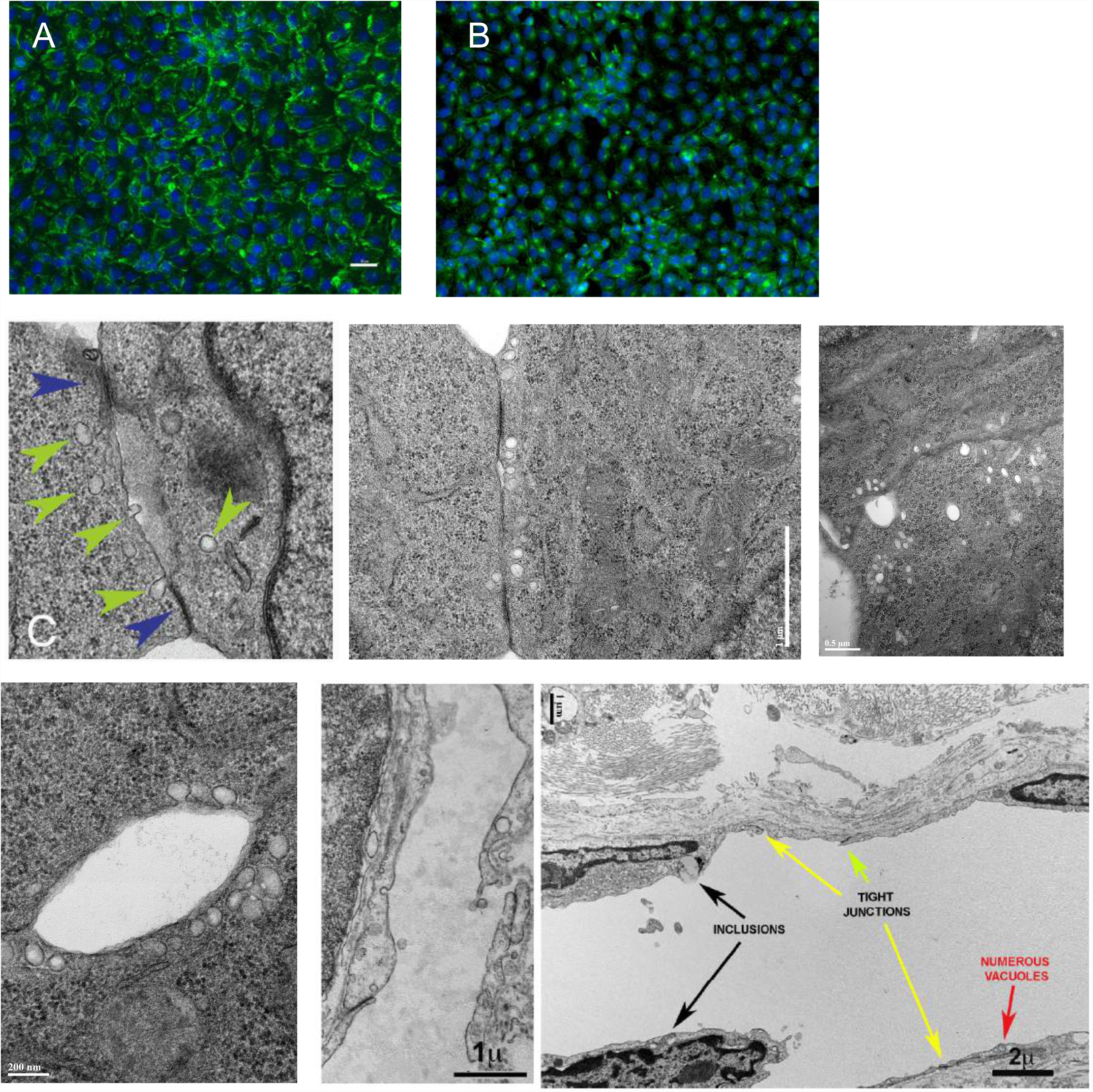
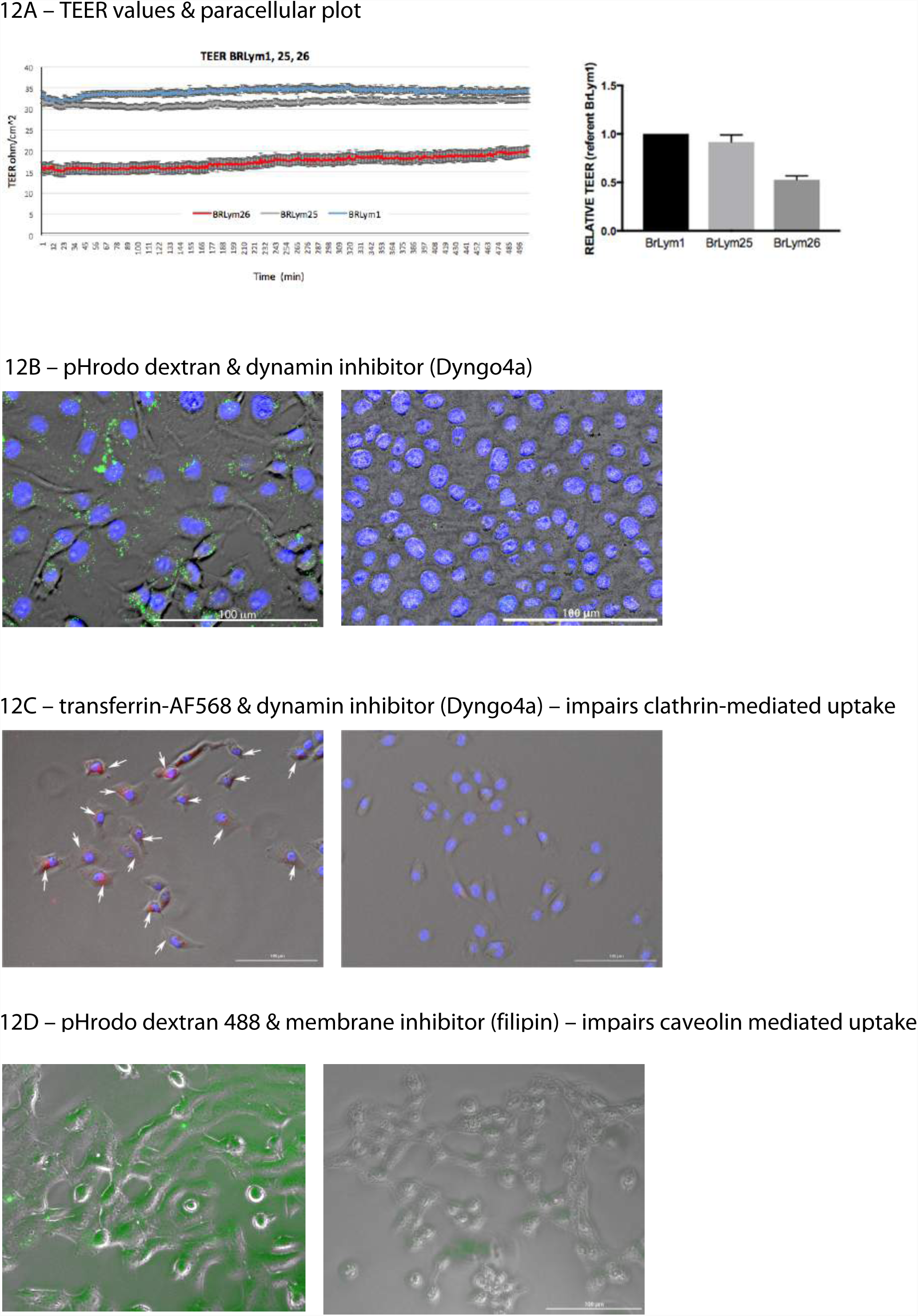
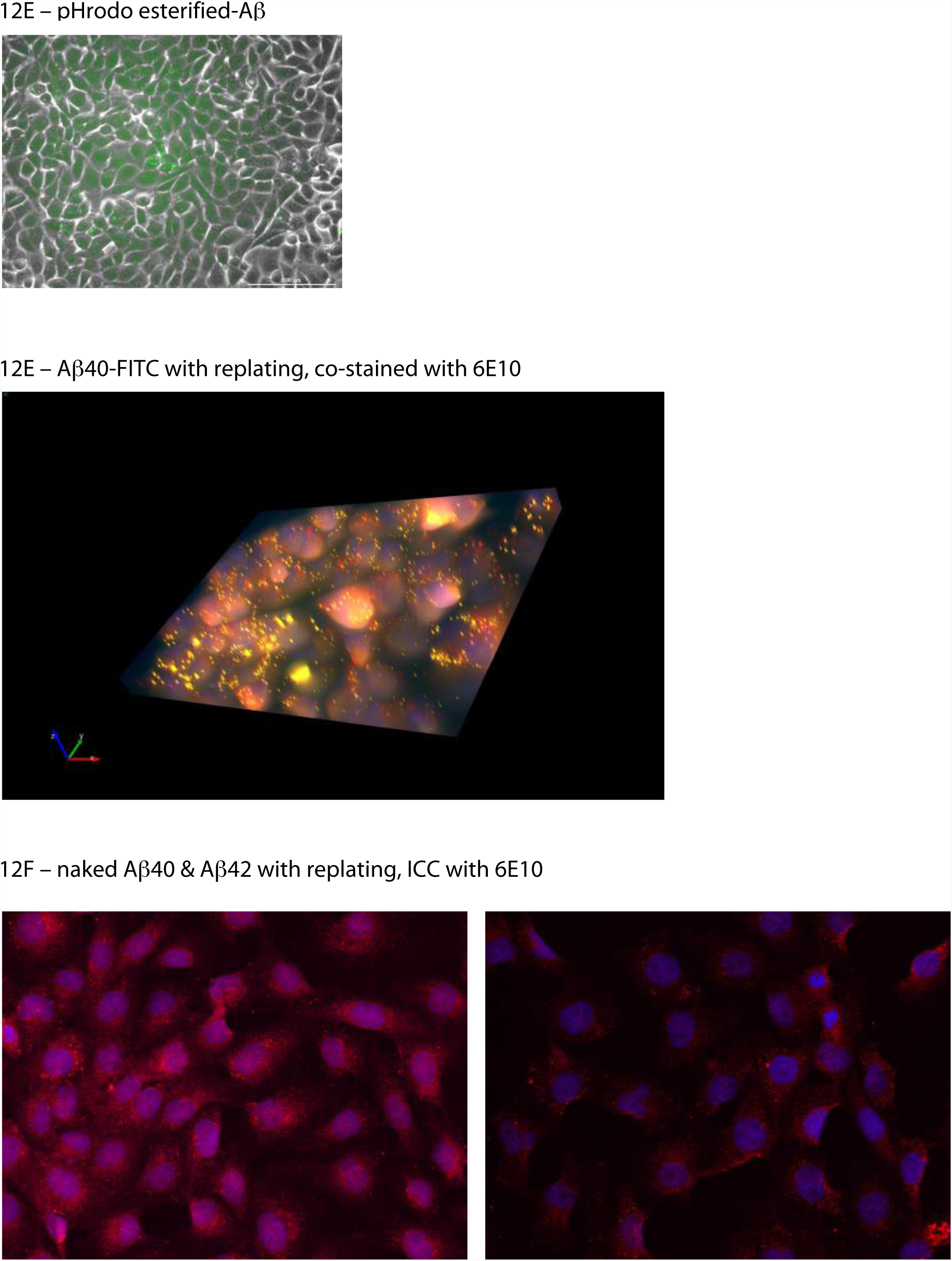
Dynamin dependent endocytosis in dural lymphatics is one mechanism of Aβ uptake.

To quantify the extent of paracellular transport *in vitro,* we performed tracer experiments in lymphatic monolayers *in vitro* using labeled dextrans of different sizes. We began by measuring the relative tightness of the membrane using continuous transepithelial electrical resistance (TEER), which is proportional to tight junction integrity. These values were uniformly higher in cell monolayers from wild-type animals compared with cells from age-matched Alzheimer’s animals, suggesting that lymphatic membranes in Alzheimer’s disease are leakier. Labeled dextran (size range 4-110kD) showed increasing permeability up to 70kD size, with a sharp drop-off thereafter. We did not detect a significant difference between wild-type and Alzheimer’s cells in transport of different sized neutral tracers in this size range. This suggests that for neutral tracers such as dextran, paracellular and transcellular transport are likely equally favored, and loss of transmembrane energy-dependent transport in Alzheimer’s cells may be partially compensated by greater membrane leakiness **[Fig 12A]**.

To isolate various mechanisms of lymphatic uptake by endocytosis, we started by performing live cell imaging using pHrodo-dextran. This is 10kD dextran conjugated to a pH sensitive dye that is not visible until internalized to endosomes. This method eliminates the possibility of detecting signal from non-specific binding to cell membranes, which can be difficult to distinguish from internalization. As expected, pHrodo dextran was readily taken up by lymphatic cells, and blocked in dose dependent fashion with Dyngo4a at 100-200μM, a specific inhibitor of dynamin-dependent endocytosis (both clathrin and caveolin mediated) **[Fig. 12B]**. This confirms that dynamin-dependent endocytosis is active in dural lymphatic cells. To test for the presence of clathrin-dependent endocytosis, we applied transferrin-AlexaFluor647, and confirmed inhibition with Dyngo4A using live cell imaging. The lymphatic cells avidly took up labeled transferrin and were again specifically blocked by the dynamin inhibitor at the same concentration **[Fig. 12C]**. To confirm the presence of caveolin mediated uptake, we treated cells with pHrodo dextran and blocked uptake with filipin, which preferentially blocks caveolin mediated uptake **[12D]**.

Having shown that caveolin and clathrin mediated endocytosis are active in dural lymphatics, we moved on to testing Aβ uptake by lymphatics. We initially treated cells with biotinylated pHrodo + strepavidin-Aβ, but this method gave high background due to excess pHrodo. To improve signal/noise, we utilized pHrodo-esterified-Aβ and purified the chemically linked pHrodo compound using column chromatography. The treated cells showed evidence of diffuse uptake of Aβ **[Fig. 12E]**.

Because these bulky constructs are structurally different from amyloid uptake which occurs *in vivo,* we next developed a technique to treat cells with FITC-labeled Aβ40/42 and measured uptake inside the cells. Following treatment with human FITC-Aβ40/42, cells were then trypsinized, washed, and replated. Cells which retained signal had internalized the peptide. In order to control for the possible effect of the FITC label, we also treated cells with naked human Aβ40/42 and trypsinized, washed, replated, and fixed the rat cells for ICC with human-specific antibody 6E10. To optimize the 3D morphology, some cells were fixed on Transwells. Images were taken every 3μ using structured illumination and Z-stacks rendered with MBF StereoInvestigator **[Fig. 12F]**. In summary, these experiments show dynamin dependent uptake of Aβ species in dural lymphatic cells and suggest that drugs which augment energy dependent endocytosis have the potential to improve clearance of toxic Aβ *in vivo.*

### RNAseq analysis shows that lymphatic energy-dependent processes are affected in aging & Alzheimer’s

Prior studies using RNAseq to analyze gene expression data for LEC and BEC have been conducted *(57,83,84).* There have been some concerns that data acquired from passaged lymphatic cells may not reflect the reality of gene expression *in vivo (84).* Given that our purified lymphatic cells were collected by cell sorting and some initial passaging is inevitable to improve yield, our results should be approached with a certain degree of caution. However, we rapidly isolated and pooled our source tissues from multiple animals of the same genotype and age, with cell isolation performed within hours of collection. We then performed minimal passaging, using lymphatic media from Cell Biologics which optimally supports *in vivo* conditions. We confirmed that genes which are ordinarily expressed at high levels in lymphatics *in vivo* (e.g., Pdpn, Lyve1) gave the expected high levels of signal in healthy isolated lymphatic cells. Perhaps more important than the absolute gene expression levels are the relative changes in expression which occur with aging and Alzheimer’s disease, and how the between-group changes correlate with the phenotypes we observed in our functional experiments. Metacore analysis indicated a number of canonical pathways which were highly differentially regulated with lymphatic cell aging, including Notch, Wnt, and Ephrin pathways. These pathways have been previously implicated in lymphatic cell development and angiogenesis in general. Multiple genes for cell-cell adhesion were also affected, including claudins, cadherins, and other junctional proteins. Of note, a number of genes which we independently found in this study as highly differentially regulated are known risk factor genes in large human GWAS studies *(85),* including ApoJ and MEF2C and Sortilin. In addition, many genes involved in mitochondrial function, glycolysis, and cellular energetics were dysregulated in TgF344-AD lymphatic cells. Although a number of endocytosis related genes such as sortilin and Rab genes were affected, caveolin and clathrin and dynamin1 were not among those genes. Numerous genes associated with leukocyte migration, activation, and inflammation were also activated in Alzheimer’s derived cells. Of note, we collected venous endothelial cells together with dural lymphatics and had intended to compare their expression, but found that BEC appeared much more sensitive to even brief *in vitro* culturing, and far fewer BEC genes were differentially regulated. Our results are consistent with previously published data that reported upregulation of cadherin-11 and cadherin-13, sortilin, and selected ABC transporters in lymphatic vs. blood endothelial cells.

## Discussion

Over a century ago, the neurosurgeon Harvey Cushing wrote presciently: “Our knowledge of the meningeal coverings of the CNS, as well as of the part played by the fluid which circulates through and over them, has hardly kept pace with our knowledge of the nervous tissues which they envelop.” *(86)* His comments still hold true over a century later, and the Alzheimer’s disease field has much to gain from a better knowledge of the meninges. The modern period of research on Alzheimer’s disease began with the discovery of Aβ in the cerebral vasculature (actually the leptomeninges) by Glenner and others in the 1980s, and our work underscores the importance of this core finding. While amyloidosis in the leptomeninges with aging and Alzheimer’s has been common knowledge for decades, fewer studies have examined amyloid deposits in the vasculature of the pachymeninges *(23).*

Similarly, while cognitive changes following cervical lymphatic ligation have been known about for over 50 years, those observations predated the discovery of Aβ and were never considered in the context of Aβ clearance. In this paper, we present evidence of a causal relationship between impairment of cervical lymphatic drainage and accelerated Alzheimer’s disease pathology in multiple brain regions including the CSF-blood barrier. We propose that dural lymphatics are necessary for brain Aβ clearance, and appear to work together with the blood-brain barrier and blood-CSF barrier. A corollary is that the blood-brain barrier is insufficient to clear pE3-Aβ and Aβ42 species from the brain, though it appears better at clearing Aβ40. The unique role of pE3-Aβ in the context of lymphatic drainage clearly warrants further exploration in the broader context of the lymphatic system and aging, as well as the probable role of bystander cells such as the numerous Mato cells which we observed around meningeal vessels.

Recent work by other investigators has helped to show the relevance of dural lymphatic vessels (and the meninges more generally) for the Alzheimer’s field *(19,67),* but many unanswered questions remain and a number of conflicting points need to be reconciled. For example, we found that cervical lymphatic ligation causes potentiation of Aβ buildup in meninges and brain. However, unlike DaMesquita *et al.,* we found diffuse Aβ throughout the meninges in non-ligated TgF344-AD animals, and also detected low amounts in wild-type animals using both IHC and biochemical techniques. In an aged wild-type animal ligated for a particularly long period (6 months), we found a surprising buildup of diffuse meningeal Aβ which was detected by 4G8 monoclonal antibody. Hence, the first-order term in modeling clearance is likely to be the length of lymphatic dysfunction, rather than age per se, and this relative risk is modified by Alzheimer’s risk factors which increase Aβ production. It is also likely that Aβ is a dynamic process that depends on a variety of other microvascular structures including meningeal arteries and arterioles.

We propose that commonly observed changing ratios of Aβ in CSF over the course of Alzheimer’s disease (i.e., gradual decrease in abeta42/40 ratio) may occur due to lymphatic failure with aging, leading to Aβ42 and pE3-Aβ retention in the brain and meninges, with greater fibrillar accumulation. This process may outpace the deterioration or the overall effect size of blood-brain barrier failure with aging. Consistent with their function as waste efflux pathways, dural lymphatics and CSF-blood barrier structures both accumulate Aβ at early time points, presaging buildup in neurons and later accumulations of tau in cognitively important brain regions. Experimental blockage of this lymphatic drainage potentiates Alzheimer’s disease pathology in multiple brain regions critical for learning and memory.

A unique feature of lymphatics appears to be their preferential uptake of pE3-Aβ, possibly due in part to in situ conversion of Aβ through upregulated glutamyl cyclase. The reason for this differential activity of glutamyl cyclase in lymphatics is unclear, or if it confers any advantage to clearance. This important question will be explored in our future work. The fact that pE3-Aβ is implicated as a particularly virulent form of Aβ which exists in the cores of plaques and precedes widespread amyloidosis and tau accumulation makes it a promising target in the context of clearance by dural lymphatics, arachnoid membranes, and other meningeal vasculature. Based on our data, we propose a new model of Aβ clearance. In this model, Aβ and pE3-Aβ begin to accumulate during aging in the hippocampus and other brain areas. Paravascular flow directs these peptides to the interstitial fluid around dural sinuses and to the CSF. In the latter compartment, Aβ is transported by arachnoid cells into the dural interstitial space and directly into the calvarium. In these locations, lymphatic vessels and veins clear pE3-Aβ and Aβ in concert with perivascular immune cells.

Much of the work in the Alzheimer’s field is based upon use of transgenic mice, and yet very few mouse models accurately replicate the natural history of Alzheimer’s disease with respect to pE3-Aβ. Timing of accumulation of pE3-Aβ varies greatly among mouse models, with some expressing it much earlier than others. Transgenic mice which produce higher levels of pE3-Aβ have more severe neuronal loss. It has been reported that 5xFAD mice exhibit pE3-Aβ starting at 4 months, but relatively low levels until 12 months or later; TgSwDI mice accumulate pE3-Aβ starting at 3 months, but relatively low levels until 24 months; and Tg2576 mice accumulate very little pE3, which occurs late in the disease course at 16 months *(43). While most transgenic mouse models show pE3-A*β *accumulation only in a subset of compacted plaques and as vascular amyloid in later stages of disease, humans exhibit pE3-A*β *in diffuse and compacted plaques and intraneuronally at earlier stages.* This pivotal toxicity is associated with neurodegeneration, impairment of LTP, and cognitive decline. The TgF344-AD rat that we used for this study demonstrates accumulation of pE3-Aβ starting at 4-8 months, and is therefore an excellent model for studying the contribution of the lymphatic system to Alzheimer’s disease. In terms of mouse models, the TBA2 mouse expresses large amounts of intraneuronal pE3-Aβ and may be the best available murine model in this context. However, the behavioral repertoire of the rat as well as the complexity of its meningeal vasculature make it a highly desirable model. Moreover, our group has cross-bred theTgF344-AD rat with a Prox1-eGFP lymphatic labeled rat, which now allows us to explore this anatomy with greater ease.

The extension of our findings from rodent to human tissue is also critical, especially given past barriers to analysis of human meninges. Our demonstration that whole-mount imaging of cleared human meninges is possible will lead to further characterization of these microvascular networks, and further improvements in clearing technology are inevitable. Recent studies involving human meninges have commented on the high level of autofluorescent material in meninges *(67).* Spectral unmixing was proposed as a solution to this problem. Despite this approach, Goodman *et al.* concluded that much of their amyloid signal was non-specific, and in particular much of the signal surrounding lymphatics was presumed to be artifactual. We have also used spectral unmixing in our imaging of CSF-blood barrier tissues, but along with deconvolution and other computational approaches, this approach cannot overcome inherent limitations of the tissue, which must be addressed in other ways. Contrary to the conclusions of Goodman *et al.,* we believe that the amyloid signal within and around lymphatics is real and reflects actual clearance.

We have identified dynamin dependent endocytosis and ABC transporters as two possible mechanistic targets at the level of the lymphatic vasculature, and exploratory RNAseq analysis further implicates energy-dependent clearance pathways. The role of endocytosis and transcytosis in clearance of waste from the interstitium is not a new idea *(87,88),* nor is its potential role in Alzheimer’s disease and inflammation *(89-92).* However, the existence of these mechanisms in lymphatic cells represents a paradigm shift from the standard mechanical model of fluid and solute flux. Of note, a number of endocytosis or phagocytosis related genes have previously turned up on large human GWAS studies, such as PICALM (clathrin-mediated endocytosis), CR1/CD35 (phagocytosis), CD33 (clathrin-independent endocytosis), TREM2 (phagocytosis), BIN1 (endocytosis), SORL1 (endocytosis), EphA1 (trans-endocytosis), CLU/LRP2 (endocytosis), but a possible connection to lymphatics was not made until now.

Application of these findings to the clinic requires a much more detailed understanding of the mechanisms of clearance at the level of dural lymphatics and the CSF-blood barrier, including the diversity of vessels involved. For example, the role of initial vs. collecting lymphatics remains unclear, and structural-morphological studies will need to be completed which span a broad range of ages and Alzheimer’s disease phenotypes. The manner in which the intrinsic brain vasculature and the meningeal vasculature work together to clear pE3-Aβ and Aβ requires further study. Given its early involvement and large impact on the pathogenesis of Alzheimer’s disease, it is possible that remediating the dural lymphatic system, or potentiating it in some way, could delay the pE3-Aβ clearance failure which we observe when it is compromised. If anatomical connections with diploic spaces are confirmed, there may be less invasive ways to target the lymphatic system in the scalp and neck, which could include lymphatic massage, revascularization procedures, relatively non-invasive gene therapy approaches, or other clinical interventions. For example, subgaleal delivery of growth factors or other biologics could help to maintain proper meningeal function with aging. In this manner, we may finally achieve Harvey Cushing’s goal of understanding the full impact of the meninges on human brain disease.

## Materials and Methods

### Statistical approach

A series of statistical analyses were performed examining total Aβ (soluble + insoluble fractions) separately for pyroglutamate-Aβ, Aβ40, Aβ42, and Aβ40+Aβ42. We compared differences in mean Aβ by region (five groups), fitting repeated measures two-way analysis of variance (ANOVA) models for the null hypothesis of no difference in mean abeta between regions. We also compared mean total Aβ between treated (ligated) and untreated (non-ligated) specimens by specific subgroupings. Using multivariable generalized linear models (GLM), we regressed total Aβ on treatment, age at tissue collection, and time since ligation (months, continuous), accounting for correlation by region *(93).* We estimated adjusted coefficients with robust standard errors *(94)* and 95% confidence intervals to determine the statistical significance of observed mean differences associated with ligation. Similarly, we compared the area fraction for Aβ sampled from the hippocampus in ligated and non-ligated animals by section (Aβ 4G8 and Aβ4G8 + pE3 Aβ area fraction) using modified GLM models regressing area fraction on ligation in age-matched samples with clustering by animal-level measurements. Postestimation statistics, Akaike and Bayesian information criterion, were used to assess model fit when adding terms to individual GLM regressions. We performed sensitivity analyses varying the assumption of a Gaussian distribution for our GLM modeling with plausible alternative (inverse Gaussian and Poisson) distributions; our results were robust to specification of models under these conditions with respect to precision and statistical significance. To account for multiple statistical comparisons resulting from 15 *a priori* fit models, we calculated post hoc adjustments of the P value to maintain a conservative family-wise type I error rate of 0.05 following the approach of Bonferroni *(95).* Similarly, *in vitro* transport data from Transwell experiments were modeled with log link regression with GLM to account for multiple observations occurring at each time point, and used to model the transport graphs and to calculate P values for each condition.

### Tissue Extract Preparation

Following microdissection of fresh tissues on ice, tissue weights were recorded. Large tissues were homogenized in Dounce homogenizers and small tissues were homogenized with Triple-Pure High Impact Zirconium Beads using BeadBug homogenizer in ice-cold Cell Signaling Technology Lysis Buffer supplemented with protein inhibitors. Samples were centrifuged at 100,000×g for 1 hr at 4°C. Supernatants were collected as detergent soluble fraction. Pellets were washed and resuspended in 5M GuHCl followed by rotation at ambient temperature for 1 hr. Detergent insoluble fractions were collected after centrifugation at 16,000×g for 30 min. All fractions were aliquoted and frozen until analysis.

### Biochemical assays for pE3-Aβ, Aβ40/42 quantification

pE3-Aβ ELISA (IBL-America) and MesoScale Discovery electrochemiluminescence (MSD-ECL) multiplex V-PLEX assays were used for quantification of Aβ species. Peptides were quantified in detergent soluble and insoluble fractions according to the manufacturer’s instructions. Peptide amounts were normalized to tissue weight.

### Immunohistochemical analyses in rat and human

Due to the thickness of rat and human meninges, it was necessary to develop new techniques to isolate, clear, and mount these tissues. Following fixation with 4% PFA and blocking with wash buffer, IHC was performed to rat or human tissue with the following panel of antibodies: pan-Aβ (Biolegend, Cat.# 800709,), pE3-Aβ (Cell Signaling Technologies, Cat.#14975; Biolegend, Cat.#822301), Pdpn (Biorybt, Cat.#orb76777; Novus, Cat.#NBP2-54347R); Prox1 (Novus, Cat.#NBP1-30045AF647; Abcam Cat.#ab38692), Lyve1 (R&D, Cat.#MAB20892; Novus, Cat.#FAB20892R; Novus, Cat.#NB100-725), CD31/PECAM (R&D, Cat.#AF3628; Novus, Cat.#NB100-64796V; Novus, Cat.#NB600-562AF488; Novus, Cat.#NB100-2284; R&D, Cat.#af806), QPCT (Novus, Cat.#NBP1-81838; LSBio, Cat. #LS-C334897) tau (Abcam, Cat. #ab85055), phospho-tau (Thermo, Cat.#MN1020), PHF (Abcam, Cat.# #ab80579). Typical incubation conditions were 1:100-1:150 over 48-72 hours. After washing, tissues were mounted with custom silicone gaskets on large-format high refractive index slides and cover slipped. FocusClear and FocusDeep (CelExplorer Labs) were used to clear tissues per manufacturer’s recommendations. For quantitative analysis of rat meninges, a standardized region was acquired in all samples under identical imaging conditions, and intensity values were derived using ImageJ.

### Stereology of rat hippocampal plaques

Whole hippocampi were microdissected post-mortem on ice and immersion fixed in 4% PFA at 4°C overnight, then cryopreserved in 30% buffered sucrose. Tissues were flash frozen in OCT compound and sections sliced lengthwise into 50μm sections with a Leica SM2010R freezing sliding microtome, starting from the lateral side and including CA1, CA3, and dentate gyrus. From approximately 100 total sections per hippocampus, 7 were selected in systematic random manner for IHC from each animal, such that there was 300 μm between each section. We stained simultaneously to total Aβ (4G8, Biolegend) and pyroglutamate-Aβ (pE3, Cell Signaling Technologies). Additional staining was performed to tau (Abcam, Cat.#ab85055;), IBA1 (Abcam, Cat.#ab107159), 200kD NF heavy chain (Abcam, Cat.#ab3966). Imaging was acquired at 10x magnification (0.3 NA objective) using Zeiss AxioImager M2 fluorescence microscope with Apotome. A virtual slide for every section of tissue was created using “Slide Scan” function on MBF StereoInvestigator, with Z-stack image acquisition every 3μm. Unbiased stereology with area fraction fractionator probe was used to determine the area fraction of Aβ plaques within the hippocampus in the regions sampled, using the maximal projection view. Counting frames of 100μm^2^ were randomly applied to sample 15% of the tissue surface area. Within each counting frame are regularly spaced (5μm) crosses (+). If a cross co-localized with immunoreactive Aβ (4G8), pyroglutamate Aβ (pE3), or brain parenchyma, the cross was marked with a unique marker. Area fraction was calculated in MBF StereoInvestigator. We sampled total volumes of 14.75 mm^3^ in ligated and 17.68mm^3^ in non-ligated animals. The average C.E. was 0.02, calculated by Gunderson technique. Mean counts for ligated and unligated animals were compared using a two-tailed T-test and graphed with GraphPad Prizm.

### Image acquisition, processing, and analysis

Confocal images were aquired using Caliber RS-G4 mosaic scanning confocal and Zeiss LSM710 confocal with 10-40x long working distance objectives (0.3-0.6NA). Post-processing was performed with Zeiss deconvolution software and with Bitplane Imaris. Images were stored as digital files on a dedicated server.

### Transmission electron microscopy (TEM)

Tissues for TEM are fixed in 2% glutaraldehyde + 4% PFA, thenwashed in 0.1M Sorensen’s sodium phosphate buffer and post-fixed in buffered 1% osmium tetroxide x 1 hour. After several buffer washes, samples were dehydrated in ethanol washes ascending to 100%, followed by two changes in propylene oxide (PO) transition fluid. Preparation of cells on Transwell membranes were performed per Corning Costar recommendations for TEM. Specimens were infiltrated overnight in a 1:1 mixture of PO and LX-112 epoxy resin, and 2 hours in 100% pure LX-112 resin, and then placed in a 60C degree oven to polymerize for 3 days. Semi-thin sections (0.5-1.0 um) were cut and stained with 1% Toluidine blue-O to confirm the areas of interest via light microscopy. Ultra-thin sections (70-80 nm) were cut using a Leica ultramicrotome, collected onto 200-mesh copper grids and contrasted with 6% uranyl acetate and Reynolds lead citrate stains. Specimens were examined using a JEOL JEM-1220 transmission electron microscope operating at 80 kV. Digital micrographs were acquired using Erlangshen ES1000W Model 785 CCD camera and Digital Micrograph software.

### Animal model & ligation surgery

All surgeries were conducted under approved IACUC protocols. Microsurgical procedures were performed using the operating microscope. We developed new methods to completely ligate both the superficial and deep cervical lymphatic chains. Following anesthesia, the lymphatic chains were identified using methylene blue subdermal injection and needle injections directly into deep lymphatics. This allowed the exposure and atraumatic blockage of cervical lymphatic chains with microsurgical titanium microclips.

### RNAseq analysis

Pooled cells were collected per approved IACUC protocol from wild-type and TgF344-AD rats using cell sorting with markers to CD31 and Pdpn. These cells were minimally passaged and frozen in RNAlater solution (Thermo). Total RNA was extracted using Maxwell^®^ 16LEV simplyRNA Tissue Kit (Promega) implemented on a Maxwell16 instrument. RNA quantification was performed using a Qubit 3.0 fluorometer with RNA BR assay kit (Thermo). In addition, RNA quality was assessed using a RNA ScreenTape kit, implemented on a 2200 TapeStation System (Agilent). RNAseq libraries were prepared using Lexogen QuantSeq 3’ mRNA sequencing library preparation kit for Illumina sequencing platform-compatible libraries. This protocol provides a novel approach, as only one library fragment is produced per endogenous transcript. This can be used for more accurate determination of gene expression values, fewer total reads per sample, and shorter sequencing length. The KAPA real time PCR Library Quantification Kit was used to quantify and assess library quality. All libraries were pooled in equimolar amounts and run on an Illumina Nextseq500 instrument, using high-output kit. For RNA-seq QC and quantification, raw reads were aligned to reference genome using the BWA MEM, which efficiently maps reads with read-through into polyA tails and adapter sequences, as is common with 3’ RNAseq. We quantified expression level of genes using FeatureCounts first as raw read counts, which are suitable for differential expression analyses, and also normalized to reads-per-million for direct comparison between samples. To ensure that data used in downstream analysis was high-quality, we performed several quality-control checks: (1) the depth and quality of the raw sequencing data and the absence of sequencing artifacts was confirmed; (2) the fraction of reads aligning to the reference genome, and the fraction of alignments that are in coding sequences, was confirmed to be acceptable for accurate expression estimations; (3) the high-level clustering of samples using principle component analysis (PCA) was compared to check for biological outliers that should be removed or further investigated prior to differential analysis. Differential expression statistics (fold-change and p-value) were computed using edgeR on raw expression counts obtained from quantification. Importantly, edgeR allows us to perform multi-group and multi-factor analyses to prioritize which genes show the biggest effects overall, as well as pair-wise tests between sample conditions to determine the context of the changes as a post-hoc evaluation. In all cases we adjusted p-values for multiple testing using the false discovery rate (FDR) correction. Significant genes were determined based on FDR threshold of 10% (0.10) in the multi-group comparison.

### ABC transporter assay

Purified cells were incorporated in Enzo EFLUXX-ID Gold multidrug assay per manufacturer’s recommendations.

### Aβ uptake assays & pHrodo

Isolated cells were plated per protocol and pHrodo and dextra (Thermo) were applied per manufacturer’s recommendations. Aβ was prepared per our lab’s standard protocol for monomeric preparations using hexafluoroisopropanol (HFIP). FITC-labeled Aβ (Anaspec) was prepared per the same protocol.

### Immunohistochemistry

Hippocampal sections where prepared with a cryostat at 50mm thickness and stored in a solution of tissue antifreeze at −20C (1×PBS with 30% glycerol and 30% ethylene glycol). Prior to staining, the tissue was washed 3 times for 15 minutes in PBS. Next, tissue was incubated in a wash/block buffer composed of 0.3% Triton X-100 and 5% normal donkey serum in PBS for 2 hours at room temperature with gentle shaking. After the blocking step, primary antibodies were added at the appropriate concentration. Tissue sections incubated in primary antibody solution over-night. In the morning, sections were washed 3 times for 15min in the wash/block buffer. After the wash, secondary antibodies were added at a concentration of 1:250 diluted in wash/block buffer: goat anti-rabbit AF488 (Abcam, product #ab150081) and donkey anti-mouse AF568 (Abcam, product number ab175700). Sections incubated in secondary antibody solution for 2 hours at room temperature with gentle shaking and protected from light. Tissue was finally washed 3 times for 15 minutes in wash/block buffer, then mounted using floating sections in PBS. Sections were mounted with a DAPI containing media (Vector Laboratories, Cat. #H-1200).

### ICC

Cell were grown on glass cover slips or Transwell supports and fixed with 4% paraformaldehyde in PBS, permeabilized with 0.5% Triton X-100 in PBS for 5 min and washed before application of primary antibodies. Cells were incubated with primary antibodies for 1 hr at RT or overnight at 4°C, washed and subsequently incubated with secondary antibodies for 1 hr. Finally, cells were washed, mounted with or without prior DAPI staining.

### Transwell based permeability assays

For transport studies, cells were plated on PET Transwell inserts in 24-well format with 0.4 μ pore size or on HTS 96-well PET Transwells with 1 μ pores (Corning). Cells were cultured for 5-7 days to form cellular barrier. Culture media in Transwell plates was removed and pre-warmed clear HBSS was added into both chambers. Dextran tracers or Ab peptides were added to basal or apical chamber depending on experiments. Ab peptides were prepared as previously described. After application of a tracer, plates were incubated at 37°C and samples were taken from both apical and basal compartments in each well at defined time points and replaced with an equal volume of respective buffer. Alternatively, HTS Transwell plate was read as one unit at the endpoint of the assay. The amount of tracer crossing the monolayer was reported as permeability (cm/sec). The cleared volume was calculated by dividing the concentration in the receiver compartment by the product concentration in the donor compartment at each time point. Average cumulative volume cleared was plotted versus time and the slope was estimated by linear regression to give the mean and the standard error of the estimate. The apparent permeability was estimated based on the equation Papp = (dC/dt)/(C0 x A), where dC/dt is the slope of concentration in basal Transwell chamber as a function of time obtained directly from measurements, C0 is the initial concentration on the inlet side of the diffusion chamber, and A is area.

### Transepithelial electrical resistance (TEER) measurements

TEER readings were performed on the cell cultures prior to tracer permeability experiments. Cells were plated on PET Transwell inserts in 24-well format with 0.4 μ pore and grown as described above. A day before the tracer assay, TEER was continuously recorded overnight with ECIS instrument (Applied Biophysics, Troy, NY). Transwell inserts were transferred into ECIS 8W TransFilter adaptor and placed into CO2 incubator connected to ECIS instrument. TEER value of a blank membrane with pore size 0.4μm was subtracted from apparent TEER values of membranes with the cells to obtain the effective TEER value.

### Size differential transport experiments

Cells were seeded on 12-well Transwell plates as described above. TEER measurements were taken prior to the assay. Fluorescently labeled dextran tracers were obtained from Molecular Probes (Eugene, Oregon): 3 kDa Cascade Blue conjugated dextran, 40 kDa Tetramethylrhodamine (TMR) conjugated, 70 kDa Texas Red conjugated and FITC-150 kDa dextran. Dextran stocks were diluted to the working concentration of 20 mg/ml and placed on the apical or basal side of the wells. Alternative chamber was filled with the media without dextran was. Plates were incubated at 37°C with gentle agitation and samples were taken from target compartments at 0, 30, 60, 90 and 120 minutes and replaced with an equal volume of the media. Fluorescence of collected samples was read with BioTek microplate reader at the following settings: 375/420 nm for Cascade Blue; FITC 494/518 nm; 555/580 for TMR; and 595/613 for Texas Red.

## Supplementary Materials

Fig. S1. First discovery of human meningeal lymphatics

Fig. S2. Schematic of rat lymphatic drainage: continuity of intracranial and extracranial vessels

Fig. S3. Lymphatic ligated TgF344-AD rat shows pE3-Aβ and Aβ within intraosseous channels

Fig. S4. Surgical ligation of superficial and deep cervical lymphatics with titanium microclips

Fig. S5. TEM of normal and Alzheimer’s derived lymphatic cells *in vitro*

Fig. S6. RNAseq tables of upregulated and downregulated genes & pathway analysis

## Acknowledgements

Thanks to Dr. Paola Leone and Dr. Andrew Freese for making this work possible.

## Funding

NIH K01 AG062789-01 to L.R. and Pilot Grants from The Silver Foundation and Global Lyme Alliance to C.J.

## Author contributions

C.J. and L.R. conceived experiments and wrote the paper; H.P., K.H., D.P., O.L., D.P., J.S., E.H. provided technical support; G.C. and C.L. participated in data analysis.

## Supplemental Data

**Fig. S1.**
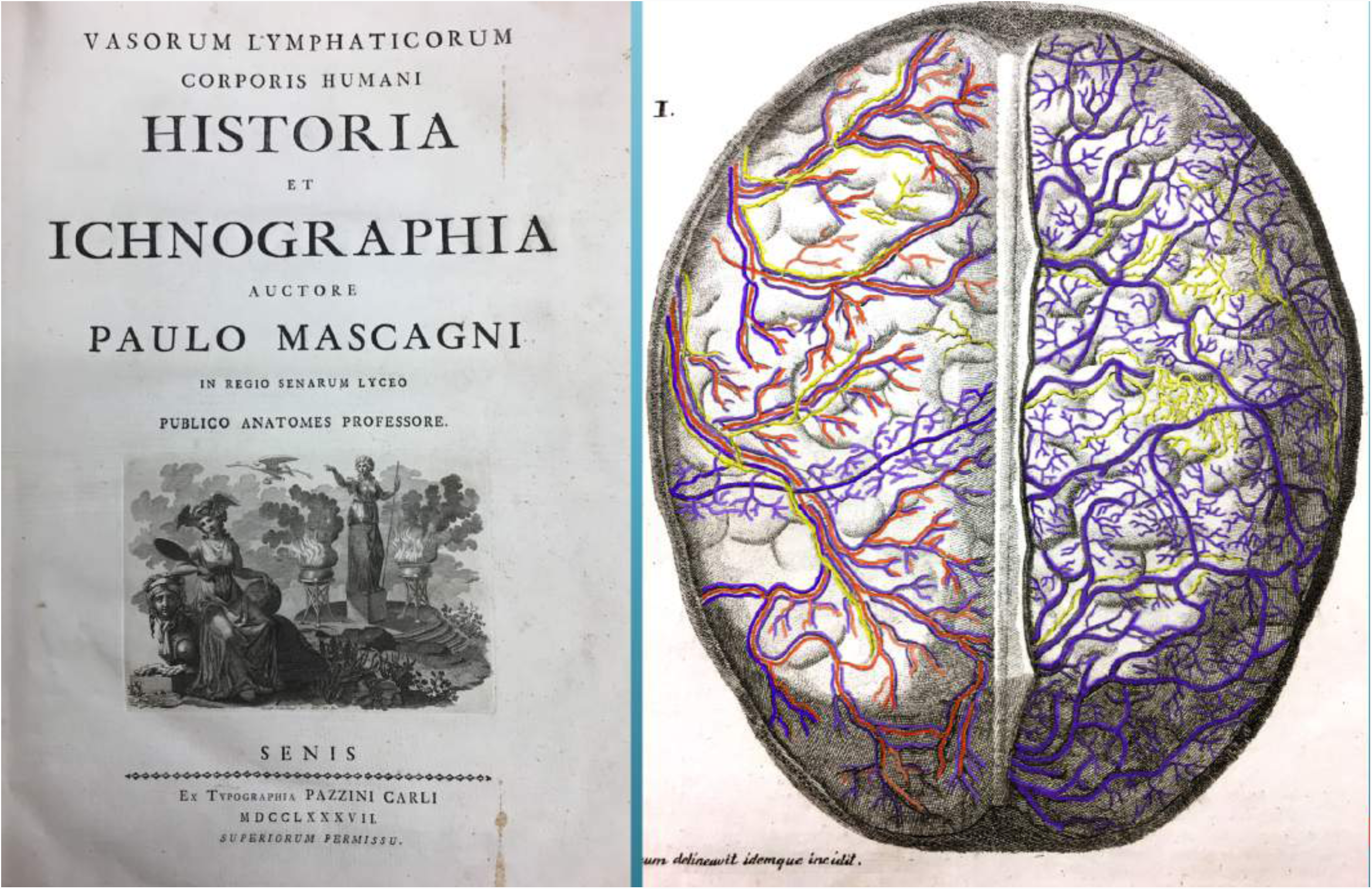
First discovery of human meningeal lymphatics. The anatomist Paolo Mascagni first demonstrated lymphatic vessels in the human meninges in his illustrated tome “Lymphatic Vessels of the Human Body” (Sienna 1787), using the mercury injection technique of his predecessor Anton Nuck (Leiden 1692). He collected specimens post-mortem and meticulously injected mercury compounds with glass microcapillary tubes (“drawn glass”). These specimens were rendered into drawings and lithography plates. To highlight the relevant anatomy, color has been added to a facsimile of his engraving of the cerebral convexity below (yellow lymphatics, blue veins, red arteries). The pattern of lymphatic collecting ducts running with meningeal vessels in the outer layers of dura (left) is consistent with what we have observed in rat and human tissue. Until now, the pattern in deeper meningeal layers has never been mapped using modern techniques of light microscopy, due to technical barriers of immunostaining and imaging this thick, opaque tissue. However, our whole-mounts of cleared human meninges imaged with fast confocal bear a striking resemblence to this depiction. While lymphatics have not been unequivocably observed in arachnoid or pia in modern preparations, Mascagni’s text suggests that lymphatic-like vessels may occupy the deeper dural and arachnoid layers after the outer layer of dura is dissected away (right).

**Fig. S2.**
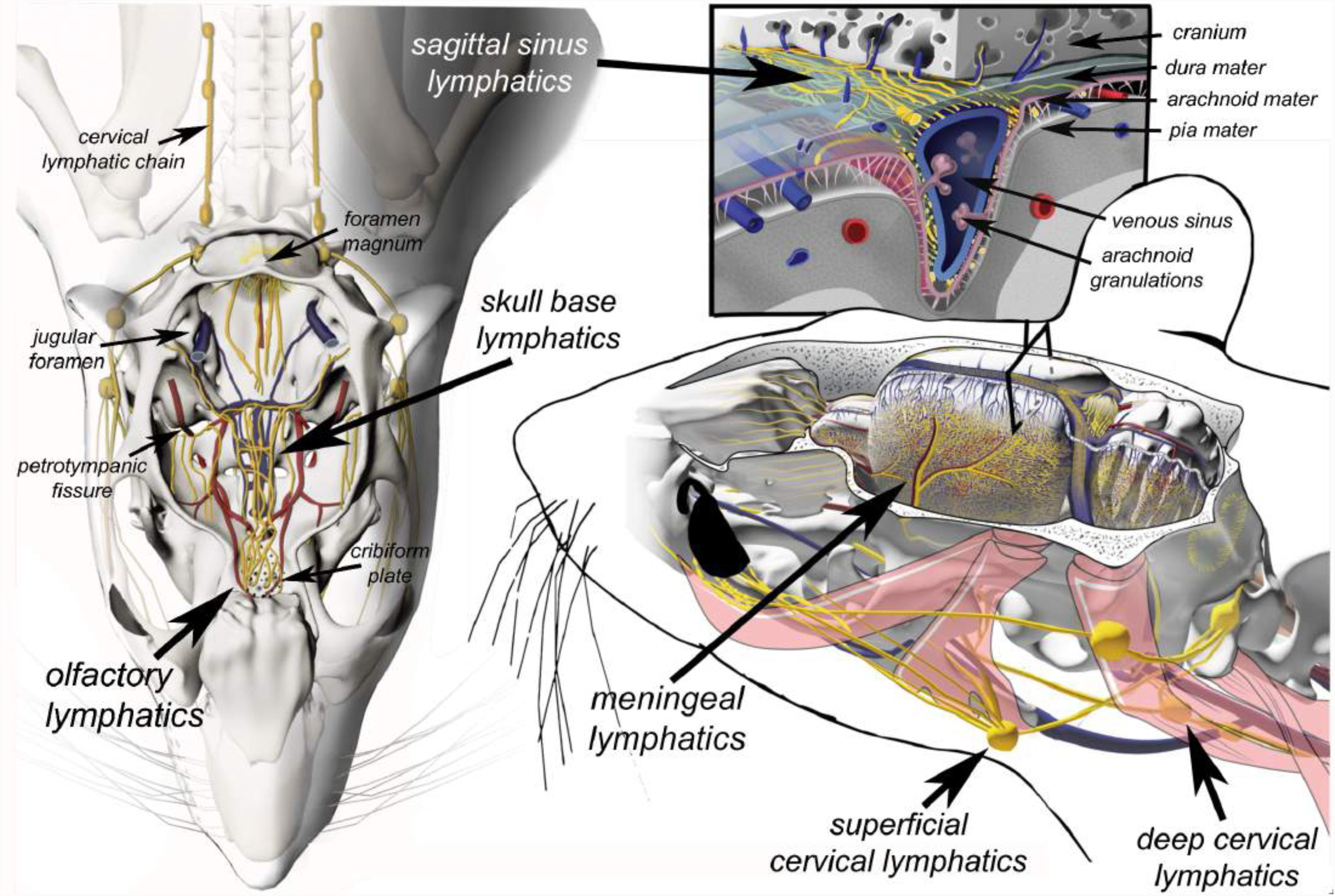
Schematic of rat lymphatic drainage: continuity of intracranial and extracranial vessels. Prior work in rodents has demonstrated dural lymphatic vessels in at least three distinct topographical areas: (1) running along meningeal artery and veins in the lateral meninges; (2) running parallel with central venous sinuses; and (3) running along the cribiform plate *(18,47,48,96).* Lymphatic collecting ducts accompanying the meningeal artery presumably exit the skull base through a plexus at the foramen spinosum. Lymphatics accompanying the central venous sinuses presumably exit the skull base through the jugular foramena. Additional lymphatics descend from the olfactory bulbs to pass through the cribiform plate. Though less well documented, lymphatic channels at the cerebral convexity may pass superiorly through venous lacunae in the skull to ramify with pericranial and scalp lymphatics. The complex relationships between meninges, vasculature, and brain are appreciated in the schematic diagram below. This computer assisted design was created by injecting the rat cervical lymphatic chain with a radiopaque tracer and imaging the head and neck with high resolution CT to view the neck lymph glands in proper relation to the skull. Representative locations of meningeal lymphatics were adapted from confocal microscopy images of whole meninges collected from 12 animals. Lymphatic vessels are shown in gold, with respect to veins, arteries, and other structures. This rendering is an initial roadmap for modeling this complex anatomy, but a full understanding will require rigorous analysis of the fine structure of vessels during aging.

**Fig. S3.**
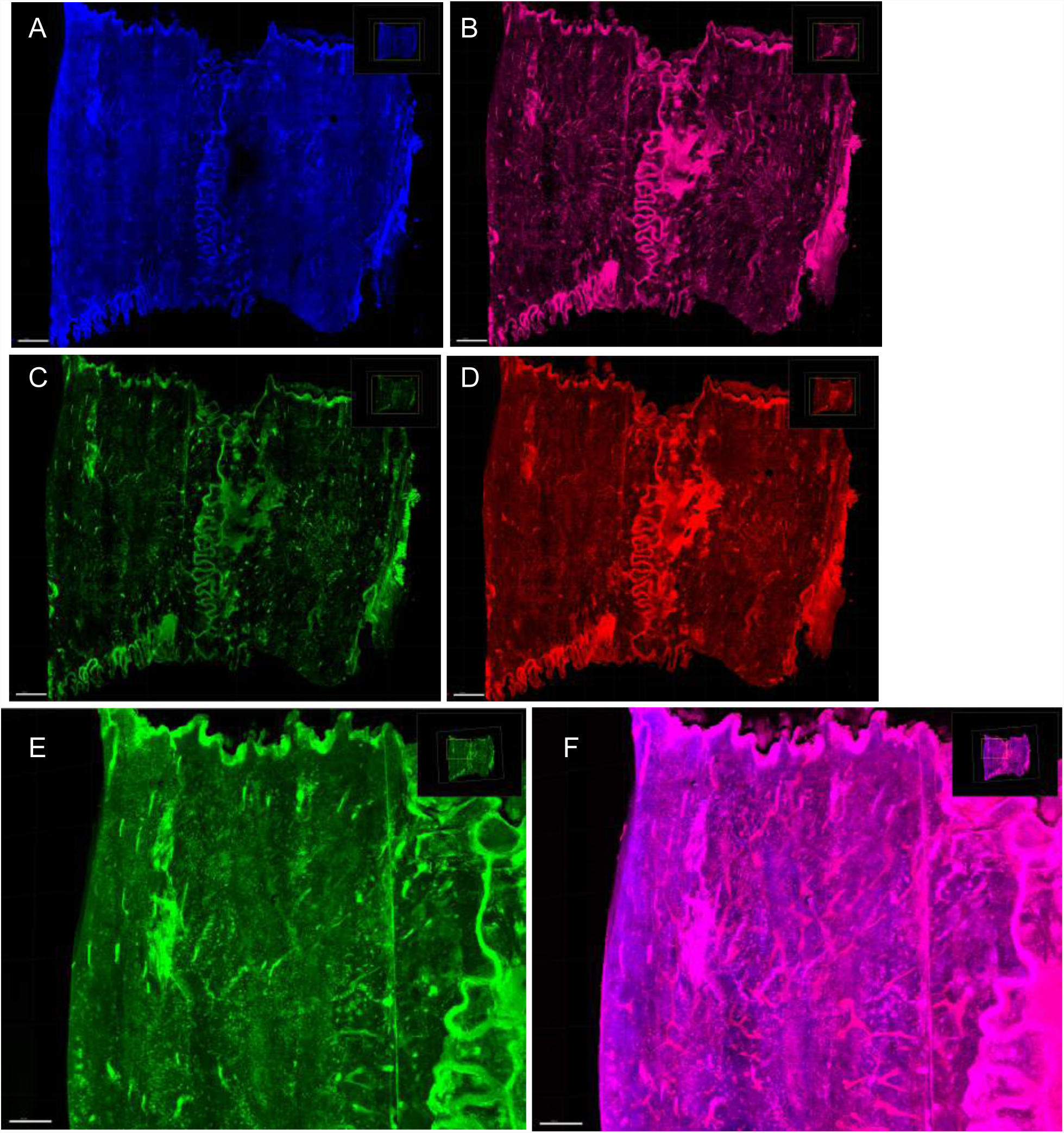
Lymphatic ligated TgF344-AD rat shows pE3-Aβ and Aβ within intraosseous channels. In addition to meninges and brain tissues, cranial bones were collected from selected animals for analysis. In this representative TgF344-AD rat, cervical lymphatics were ligated for 18 weeks (terminal sacrifice at age 9 months). The cranial vault was harvested, fixed in 4% PFA, then demineralized in 0.5M EDTA for 6 months. The clarified skull was washed, immunohistochemically stained to Aβ and vascular or lymphatic markers, and mounted in high refractive index medium. Images are maximal intensity projection, acquired every 7.62μ in Z-plane with mosaic confocal scanning microscopy, total 202 slices. Similar to the numerous immune cells around meningeal lymphatic collecting ducts and central venous sinuses in this animal, we observed numerous CD31+ Pdpn+ cells within the cleared calvarium, consistent with either macrophages or osteocytes. The majority of these cells were positive for pE3-Aβ and pan-Aβ antibody markers. We observed CD31+ Pdpn+ diploic channels in a staghorn configuration, as well as a sinuous vessel in the midline superficial to the cancellous bone. A similar configuration was previously reported in the mouse skull *(65).* The presence of strong Aβ signal in bone suggests that amyloid efflux and deposition occurs throughout the calvarium during aging. This efflux route is consistent with recent reports that arachnoid protrusions extend directly into diploic channels in bone, as well as into dural sinuses *(97,98).* Immunohistochemical markers are as follows: (A) CD31; (B) Pdpn; (C) pE3-Aβ; (D) pan-Aβ (4G8). Co-localization of pE3-Aβ and Pdpn in the midline vessel and staghorn channels is seen in (E) and (F). The precise identity of these vessels or channels is unknown, but may be lymphatic or venous.

**Fig. S4.**
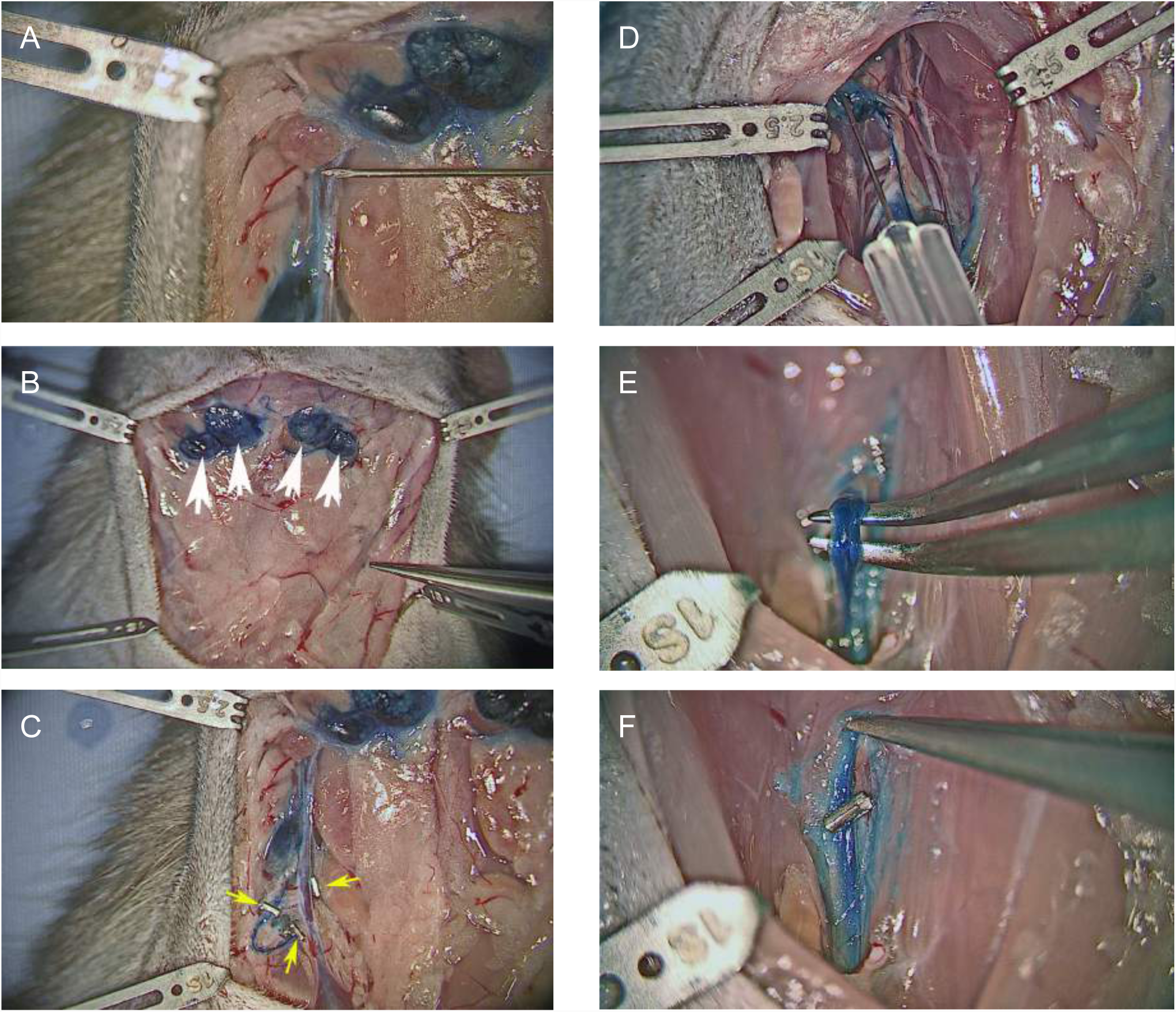
Surgical ligation of superficial and deep cervical lymphatics with titanium microclips. Superficial and deep lymphatic glands were surgically exposed and injected with methylene blue to visualize the lymphatic chains under the operating microscope, prior to permanent clipping using microclips. Unlike ablation, this blocks the cervical trunk but does not allow for regrowth. (A) Injection of superficial lymph node. (B) Four large superficial lymph nodes injected. (C) Surgical clipping of superficial chain. (D) Injection of deep cervical lymph node. (E) Mobilization and isolation of the deep cervical chain. (F) Surgical clipping of deep cervical chain; typically multiple clips are applied. Bilateral clipping of both superficial and deep chains is achieved under the operating microscope.

**Fig. S5.**
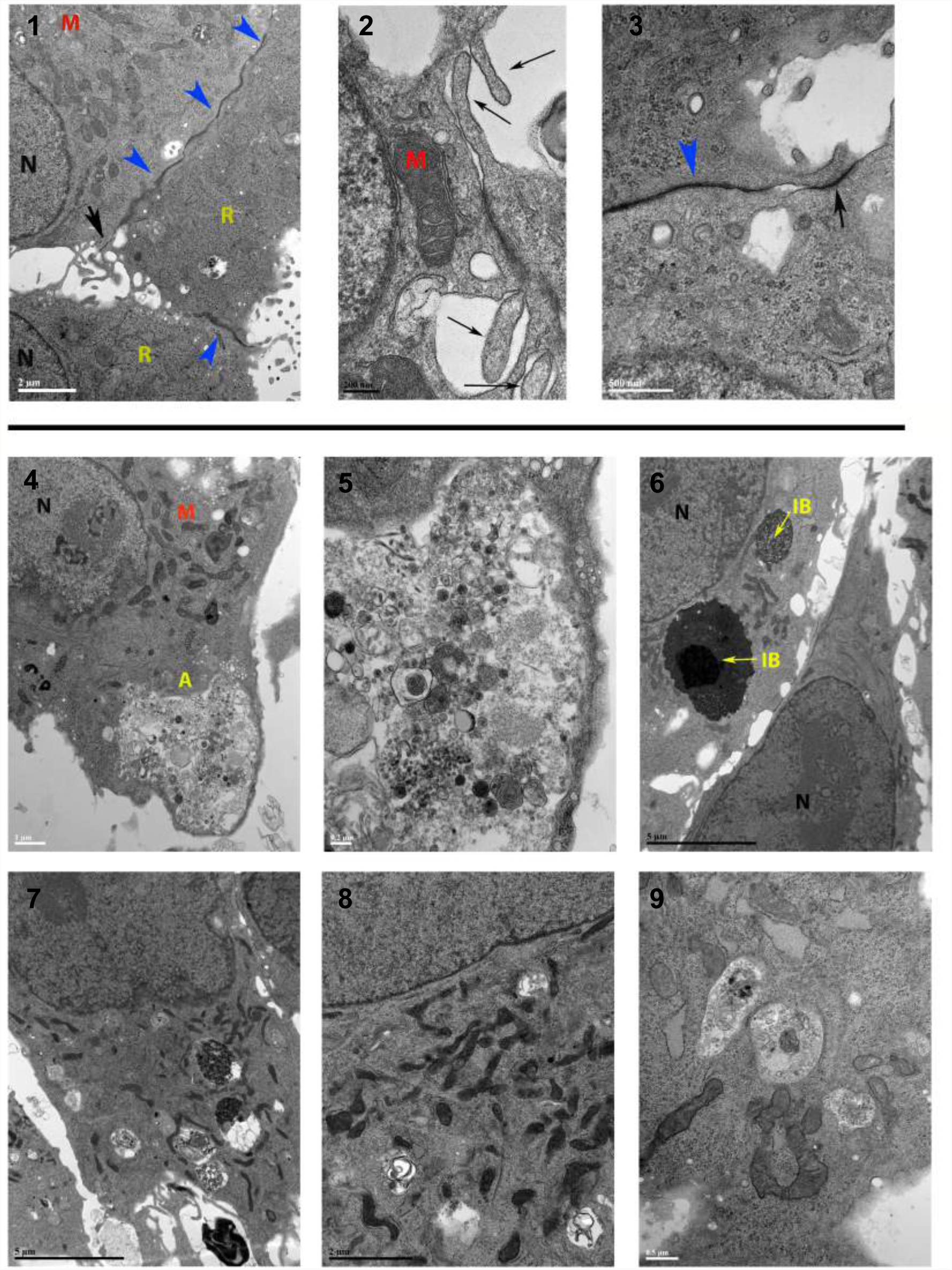
TEM of normal and Alzheimer’s derived lymphatic cells *in vitro*. **(1)** Healthy mitochondria (M), ribosomes (R), nuclei (N), and intercellular junctions (blue) in dural lymphatic cells collected from 18 month wild-type animals, magnification 17,100x. **(2)** Flap-like overleafs at intercellular junctions and healthy mitochondria (M), in dural lymphatic cells collected from 12 month wild-type animals, mag. 81,900x. **(3)** Continuous electron-dense junction (blue arrow) and tight junction (black arrow), in dural lymphatic cells collected from 18 month animals, mag. 66,900x. The remainder of cells were derived from 18 month Alzheimer’s animals. **(4)** Autophagy (A), with disrupted nuclear membrane, mag. 27,600x. **(5)** Close-up of autophagy, lipid droplets and pigmented granules, mag. 54,600x. **(6)** Pigmented intracellular inclusion bodies, mag. 11,600x. **(7)** Abnormal, sparse mitochondria and multiple cytosolic inclusions, mag. 11,600x. **(8)** Irregular mitochondria and inclusions, mag. 22,000x. **(9)** Irregular mitochondria and inclusions, mag. 33,800x.

**Fig. S6. RNAseq tables of upregulated and downregulated genes & pathway analysis**

4 months wild-type cells vs. 18 months wild-type cells 18 months wild-type cells vs. 18 months Alzheimer cells

- Pathway analysis – separately posted on web server
- List of top dysregulated genes – separately posted on web server

## References

1. H.F. Cserr, C.S. Patlak, “Secretion and bulk flow of interstitial fluid” in Physiology and Pharmacology of the Blood-Brain Barrier (Springer-Verlag, Berlin, 1992), pp. 245–261.

2. J.R. Casley-Smith, E. Foldi-Borcsok, M. Foldi, The prelymphatic pathways of the brain as revealed by cervical lymphatic obstruction and the passage of particles. Br J Exp Path 57,179–188 (1976).

3. J.J. Iliff, M. Wang, Y. Liao, B.A. Plogg, W. Peng, G.A. Gundersen, H. Benveniste, G.E. Vates, R. Deane, S.A. Goldman, E.A. Nagelhus, M. Nedergaard, A paravascular pathway facilitates CSF flow through the brain parenchyma and the clearance of interstitial solutes, including amyloid-β. Sci Transl Med (2012) 15(147), 147ra111.

4. P. Mascagni, Vasorum lymphaticorum corporis humani (Pazzini Carli, Sienna, 1787).

5. M. Foldi, J.R. Casley-Smith, Eds., Lymphangiology (Schattauer Verlag, Stuttgart, 1983), pp. 475–533.

6. L.H. Weed, The absorption of cerebrospinal fluid into the venous system. Amercian Journal of Anatomy 31(3), 191–221 (1922).

7. A.D. Speransky, A Basis for the Theory of Medicine (International Publishers, New York, 1943).

8. M. Foldi, A. Gellert, M. Kozma, M. Poberai, O.T. Zoltan, E. Csanda E, New contributions to the anatomical connections of the brain and the lymphatic system. Acta Anat 64, 498–505 (1966).

9. I. Rusznyak, M. Foldi, G. Szabo G, Lymphatics and lymph circulation, physiology and pathology. (Pergamon Press, Oxford, 1967).

10. M.W.B. Bradbury, H.F. Cserr, R.J. Westrop, Drainage of cerebral interstitial fluid into deep cervical lymph of the rabbit. Am J Physiol 240, F329–336 (1981).

11. S. Kida, A. Pantazis, R.O. Weller, Cerebrospinal fluid drains directly from the subarachnoid space into nasal lymphatics in the rat: anatomy, histology, and immunological significance. Neuropathol Appl Neurobiol 19,480–488 (1993).

12. M. Boulton, A. Young, J. Hay, D. Armstrong, M. Flessner, M. Schwartz, M. Johnston, Drainage of CSF through lymphatic pathways and arachnoid villi in sheep: measurement of 125I-albumin clearance. Neuropathol Appl Neurobiol 22(4), 325–33 (1996).

13. M.W.B. Bradbury, R.J. Westrop. “Lymphatics and the drainage of cerebrospinal fluid,” in Hydrocephalus, K. Shapiro, A. Marmarou, H. Portnoy, Eds. (Raven, New York, 1984).

14. M. Foldi, B. Csillik, O.T. Zoltan, Lymphatic drainage of the brain. Experientia, 1283–1287 (1968).

15. K.K. Ball, N.F. Cruz, R.E. Mrak, G.A. Daniel, Trafficking of glucose, lactate, and amyloid beta from the inferior coliculus through perivascular routes. J Cereb Blood Flow Metab 30,162–176 (2010).

16. M. Pappolla, K. Sambamurti, R. Vidal, Pacheco-Quinto J, Poeggeler B, Matsubara E, Evidence for lymphatic Aβ clearance in Alzheimer’s transgenic mice. Neurobiol Dis 71, 215–9 (2014).

17. A. Louveau, I. Smirnov, T.J. Keyes, J.D. Eccles, S.J. Rouhani, J.D. Peske, N.C. Derecki, D. Castle, J.W. Mandell, K.S. Lee, T.H. Harris, J. Kipnis, Structural and functional features of central nervous system lymphatic vessels. Nature 523(7560), 337–41 (2015).

18. A. Aspelund, S. Antila, S.T. Proulx, T.V. Karlsen, S. Karaman, M. Detmar, H. Wiig, K. Alitalo, A dural lymphatic vascular system that drains brain interstitial fluid and macromolecules. J Exp Med 212(7), 991–9 (2015).

19. S. DaMesquita, A. Louveau, A. Vaccari, I. Smirnov, R.C. Cornelison, K.M. Kingsmore, C. Contarino, S. Onengut-Gumuscu, E. Farber, D. Raper, K.E. Viar, R.D. Powell, W. Baker, N. Dabhi, R. Bai, R. Cao, S. Hu, S.S. Rich, J.M. Munson, M.B. Lopes, C.C. Overall, S.T. Acton, J. Kipnis, Functional aspects of meningeal lymphatics in ageing and Alzheimer’s disease. Nature 560(7717),185-191 (2018).

20. G.G. Glenner, C.W. Wong, Alzheimer’s disease: initial report of the purification and characterization of a novel cerebrovascular amyloid protein. Biochem Biophys Res Commun. 120(3), 885–90 (1984).

21. T. Hamano, M. Yoshimura, T. Yamazaki, Y. Shinkai, K. Yanagisawa, M. Kuriyama, Y. Ihara, Amyloid beta-protein accumulation in the leptomeninges during aging and in Alzheimer disease. J Neuropathol Exp Neurol. 56, 922–32 (1997).

22. Y. Shinkai, M. Yoshimura, M. Morishima-Kawashima, Y. Ito, H. Shimada, K. Yanagisawa, Y. Ihara, Amyloid beta-protein deposition in the leptomeninges and cerebral cortex. Ann Neurol. 42:899–908 (1997).

23. G.G. Kovacs, M.I. Lutz, G. Ricken, T. Ströbel, R. Höftberger, M. Preusser, G. Regelsberger, S. Hönigschnabl, A. Reiner, P. Fischer, H. Budka, J.A. Hainfellner, Dura mater is a potential source of Aβ seeds. Acta Neuropathol. 131:911–23 (2016).

24. K.G. Mawuenyega, W. Sigurdson, V. Ovod, L. Munsell, T. Kasten, J.C. Morris, K.E. Yarasheski, R.J. Bateman, Decreased clearance of CNS beta-amyloid in Alzheimer’s disease. Science 330(6012), 1774 (2010).

25. M. Foldi, E. Foldi (Eds), Foldi’s Textbook of Lymphology, 3rd Ed. Elsevier: Munich, 2012.

26. S. Jawdar, O. Wirths, T.A. Bayer, Pyroglutamate-abeta, a hatchet man in Alzheimer disease. J Biol Chem 286, 38825–38832 (2011).

27. E.K. Agare, S.R. Leonard, G.L. Curran, C.C. Yu, V.J. Lowe, A.K. Paravastu, J.F. Poduslo, K.K. Kandimalla, Traffic jam at the blood-brain barrier promates greater accumulation of Alzheimer’s disease Aβ proteins in the cerebral vasculature. Molec Pharmaceutics 10,1557–1565 (2013).

28. M.M. Verbeek, R.M.W. de Waal, H.V. Vinters. “Cerebral amyloid angiopathy” in Alzheimer’s disease and related disorders. (Springer, New York, 2000).

29. R.M. Cohen, K. Rezai-Zadeh, T.M. Weitz, A. Rentsendorj, D. Gate, I. Spivak, Y. Bholat, V. Vasilevko, C.G. Glabe, J.J. Breunig, P. Rakic, H. Davtyan, M.G. Agadjanyan, V. Kepe, J.R. Barrio, S. Bannykh, C.A. Szekely, R.N. Pechnick, T. Town, A transgenic Alzheimer rat with plaques, tau pathology, behavioral impairment, oligomeric Aβ, and frank neuronal loss. J. Neurosci. 33(15),6245–56 (2013).

30. H. Cynis, J.L. Frost, H. Crehan, C.A. Lemere, Immunotherapy targeting pyroglutamate-3Aβ: prospects and challenges. Molec Neurodegeneration 11(48),1–11 (2016).

31. C. Russo, E. Violani, S. Salis, V. Venezia, V. Dolcini, G. Damonte, U. Benatti, C. D’Arrigo, E. Patrone, P. Carlo, G. Schettini, Pyroglutamate-modified amyloid beta-peptides-AbetaN3(pE)-strongly affect cultured neuron and astrocyte survival. J. Neurochem. 82(6), 1480–9 (2002).

32. A.P. Gunn, B.X. Wong, T. Johanssen, J.C. Griffith, C.L. Masters, A.I. Bush, K.J. Barnham, J.A. Duce, R.A. Cherny, Amyloid-β Peptide Aβ3pE-42 Induces Lipid Peroxidation, Membrane Permeabilization, and Calcium Influx in Neurons. J Biol Chem. 291(12), 6134–45 (2016).

33. E.N. Cline, M.A. Bicca, K.L. Viola, W.L. Klein, The amyloid-β oligomer hypothesis, beginning of the third decade. J. Alz. Research 64, S567–610 (2018).

34. A. Piccini, C. Russo, A. Gliozzi, A. Relini, A. Vitali, R. Borghi, L. Giliberto, A. Armirotti, C. D’Arrigo, A. Bachi, A. Cattaneo, C. Canale, S. Torrassa, T.C. Saido, W. Markesbery, P. Gambetti, M. Tabaton, beta-amyloid is different in normal aging and in Alzheimer disease. J. Biol. Chem. 280(40), 34186–92 (2005).

35. J.M. Nussbaum, S. Schilling, H. Cynis, A. Silva, E. Swanson, T. Wangsanut, K. Tayler, B. Wiltgen, A. Hatami, R. Rönicke, K. Reymann, B. Hutter-Paier, A. Alexandru, W. Jagla, S. Graubner, C.G. Glabe, H.U. Demuth, G.S. Bloom, Prion-like behaviour and tau-dependent cytotoxicity of pyroglutamylated amyloid-β. Nature. 485, 61–5 (2012).

36. L. De Kimpe, E.S. van Haastert, A. Kaminari, R. Zwart, H. Rutjes, J.J. Hoozemans, W. Scheper, Intracellular accumulation of aggregated pyroglutamate amyloid beta: convergence of aging and Aβ pathology at the lysosome. Age (Dordr). 35(3), 673–87 (2013).

37. D. Schlenzig, R. Rönicke, H. Cynis, H.H. Ludwig, E. Scheel, K. Reymann, T. Saido, G. Hause, S. Schilling, H.U. Demuth, N-Terminal pyroglutamate formation of Aβ38 and Aβ40 enforces oligomer formation and potency to disrupt hippocampal long-term potentiation. J. Neurochem. 121(5), 774–84 (2012).

38. T.C. Saido, T. Iwatsubo, D.M. Mann, H. Shimada, Y. Ihara, S. Kawashima, Dominant and differential deposition of distinct beta-amyloid peptide species, A beta N3(pE), in senile plaques. Neuron 14(2), 457–66 (1995).

39. Y. Harigaya, T.C. Saido, C.B. Eckman, C.M. Prada, M. Shoji, S.G. Younkin, Amyloid beta protein starting pyroglutamate at position 3 is a major component of the amyloid deposits in the Alzheimer’s disease brain. Biochem Biophys Res Commun. 276(2), 422–7 (2000).

40. G. Wu, R.A. Miller, B. Connolly, J. Marcus, J. Renger, M.J. Savage, Pyroglutamate-modified amyloid-β protein demonstrates similar properties in an Alzheimer’s disease familial mutant knock-in mouse and Alzheimer’s disease brain. Neurodegener Dis. 14(2), 53–66 (2014).

41. S. Schilling, T. Lauber, M. Schaupp, S. Manhart, E. Scheel, G. Böhm, H.U. Demuth, On the seeding and oligomerization of pGlu-amyloid peptides (in vitro). Biochemistry 45(41), 12393–9 (2006).

42. D. Schlenzig, S. Manhart, Y. Cinar, M. Kleinschmidt, G. Hause, D. Willbold, S.A. Funke, S. Schilling, H.U. Demuth, Pyroglutamate formation influences solubility and amyloidogenicity of amyloid peptides. Biochemistry 48(29), 7072–8 (2009).

43. Frost JL, Le KX, Cynis H, Ekpo E, Kleinschmidt M, Palmour RM, Ervin FR, Snigdha S, Cotman CW, Saido TC, Vassar RJ, St George-Hyslop P, Ikezu T, Schilling S, Demuth HU, Lemere CA, Pyroglutamate-3 amyloid-β deposition in the brains of humans, non-human primates, canines, and Alzheimer disease-like transgenic mouse models. Am. J. Pathol. 183(2), 369–81 (2013).

44. O. Wirths, H. Breyhan, H. Cynis, S. Schilling, H.U. Demuth,T.A. Bayer, Intraneuronal pyroglutamate-Abeta 3-42 triggers neurodegeneration and lethal neurological deficits in a transgenic mouse model. Acta Neuropathol. 118(4), 487–96 (2009).

45. J.L. Wittnam, E. Portelius, H. Zetterberg, M.K. Gustavsson, S. Schilling, B. Koch, H.U. Demuth, K. Blennow, O. Wirths, T.A. Bayer, Pyroglutamate amyloid β (Aβ) aggravates behavioral deficits in transgenic amyloid mouse model for Alzheimer disease. J. Biol. Chem. 287(11), 8154–62 (2012).

46. A. Dorr, B. Sahota, L. Chinta, M. Brown, A. Lai, K. Ma, C.A. Hawkes, J. McLaurin, B. Stefanovic, Amyloid-β dependent compromise of microvascular structure and function in a model of Alzheimer’s disease. Brain 135, 3039–3050 (2012).

47. R.M. Izen, T. Yamazaki, Y. Nishinaka-Arai, Y.K. Hong, Y. Mukouyama, Postnatal development of lymphatic vasculature in the brain meninges. Dev. Dynamics 247(5), 741–753 (2018).

48. S. Antila, S. Karaman, H. Nurmi, M. Airavaara, M.H. Voutilainen, T. Mathivet, D. Chilov, Z. Li, T. Koppinen, J.H. Park, S. Fang, A. Aspelund, M. Saarma, A. Eichmann, J.L. Thomas, K. Alitalo K, Development and plasticity of meningeal lymphatic vessels. J Exp Med. 214(12), 3645–3667 (2017).

49. V. Shukla, L.A. Hayman, C. Ly, G. Fuller, K.H. Taber, Adult cranial dura, instrinsic vessels. J. Computer Assisted Tomography 26, 1069–1074 (2002).

50. S.J.C. Fishpool, N. Suren, F. Roncaroli, H. Ellis, Middle meningeal artery hemorrhage: an incorrect name. Clinical Anatomy 20, 371–375 (2007).

51. A.M. Kerrigan, L. Navarro-Nuñez, E. Pyz, B.A. Finney, J.A. Willment, S.P. Watson, G.D. Brown. Podoplanin-expressing inflammatory macrophages activate murine platelets via CLEC-2. J. Thromb Haemost. 10(3), 484-6 (2012)

52. M. Mato, S. Ookawara, A. Sakamoto, E. Aikawa, T. Ogawa, U. Mitsuhashi, T. Masuzawa, H. Suzuki, M. Honda, Y. Yazaki, E. Watanabe, J. Luoma, S. Yla-Herttuala, I. Fraser, S. Gordon, T. Kodam, Involvement of specific macrophage-lineage cells surrounding arterioles in barrier and scavenger function in brain cortex. PNAS 93(8), 3269–74 (1996).

53. C.A. Hawkes, J. McLauren J, Selective targeting of perivascular macrophages for clearance of beta-amyloid in cerebral amyloid angiopathy. PNAS 106(4), 1261–1266 (2009).

54. J. Zaghi, B. Goldenson, M. Inayathullah, A.S. Lossinsky, A. Masoumi, H. Avagyan, M. Mahanian, M. Bernas, M. Weinand, M.J. Rosenthal, A. Espinosa-Jeffrey, J. DeVellis, D.B. Teplow, M. Fiala, Alzheimer disease macrophages shuttle amyloid-beta from neurons to vessels, contributing to amyloid angiopathy. Acta Neuropath. 117,111–124 (2009).

55. J. Hur, J.H. Jang, I.Y. Oh, J.I. Choi, J.Y. Yun, J. Kim, Y.E. Choi, S.B. Ko, J.A. Kang, J. Kang, S.E. Lee, H. Lee, Y.B. Park, H.S. Kim, Human podoplanin-positive monocytes and platelets enhance lymphangiogenesis through the activation of the podoplanin/CLEC-2 axis. Mol Ther. 22(8), 1518–1529 (2014).

56. M. Morawski, S. Schilling, M. Kreuzberger, A. Waniek, C. Jäger, B. Koch, H. Cynis, A. Kehlen, T. Arendt, M. Hartlage-Rübsamen, H.U. Demuth, S. Roßner S, Glutaminyl cyclase in human cortex: correlation with (pGlu)-amyloid-β load and cognitive decline in Alzheimer’s disease. J. Alzheimers Dis. 39(2), 385–400 (2014).

57. C. Bridel, T. Hoffmann, A. Meyer, S. Durieux, M.A. Koel-Simmelink, M. Orth, P. Scheltens, I. Lues, C.E. Teunissen, Glutaminyl cyclase activity correlates with levels of Aβ peptides and mediators of angiogenesis in cerebrospinal fluid of Alzheimer’s disease patients. Alzheimers Res Ther. 9(1), 38 (2017).

58. G.M. Nelson, T.P. Padera, I. Garkavtsev, T. Shioda, Jain RK, Differential gene expression of primary cultured lymphatic and blood vascular endothelial cells. Neoplasia 9(12), 1038–45 (2007).

59. S. Schilling, U. Zeitschel, T. Hoffmann, U. Heiser, M. Francke, A. Kehlen, M. Holzer, B. Hutter-Paier, M. Prokesch, M. Windisch, W. Jagla, D. Schlenzig, C. Lindner, T. Rudolph, G. Reuter, H. Cynis, D. Montag, H.U. Demuth, S. Rossner, Glutaminyl cyclase inhibition attenuates pyroglutamate Abeta and Alzheimer’s disease-like pathology. Nat. Med. 14(10), 1106–11 (2008).

60. A. Azaripour, T. Lagerweij, C. Scharfbillig, A.E. Jadczak, B. Willershausen, C.J.F. Van Noorden, A survey of clearing techniques for 3D imaging of tissues with special reference to connective tissue, Prog. Histochem. Cytochem. 51, 9–23 (2016).

61. M. Ke, S. Fujimoto, T. Imai, SeeDB: a simple and morphology-preserving optical clearing agent for neuronal circuit reconstruction. Nat. Neuroscience 16(8), 1154–1161 (2013).

62. H. Hama, H. Hioki, K. Namiki, T. Hoshida, H. Kurokawa, F. Ishidate, T. Kaneko, T. Akagi, T. Saito, T. Saido, A. Miyawaki, ScaleS: an optical clearing palette for biological imaging. Nat Neuroscience 18(10), 1518–1529 (2015).

63. B. Hou, D. Zhang, S. Zhao, M. Wei, Z. Yang, S. Wang, J. Wang, X. Zhang, B. Liu, L. Fan, Y. Li, Z. Qiu, C. Zhang, T. Jiang, Scalable and Dil-compatible optical clearance of the mammalian brain. Front. Neuroanat. 9, 1–11 (2015).

64. L. Chen, G. Li, Y. Li, H. Zhu, L. Tang, P. French, J. McGinty, S. Ruan, UbasM: an effective balanced optical clearing method for intact biomedical imaging. Nature Reports 7, 12218 (2017).

65. S. Calve, A. Ready, C. Huppenbauer, R. Main, C.P. Neu, Optical clearing in dense connective tissues to visualize cellular connectivity in situ. PlosOne 10(1), e0116662 (2015).

66. Y. Liu, A. Chiang, High resolution confocal imaging and three-dimensional rendering. Methods 30, 86–93 (2003).

67. J.R. Goodman, Z.O. Adham, R.L. Woltjer, A.W. Lund, J.J. Iliff, Characterization of dural sinus-associated lymphatic vasculature in human Alzheimer’s dementia subjects. Brain, Behavior, Immunity 73, 34–40 (2018).

68. D. Predescu, R. Horvat, S. Predescu, G. Pallade, Transcytosis in the continuous endothelium of the myocardial microvasculature is inhibited by N-ethylmaleimide. PNAS 91, 3014–2018 (1994).

69. D. Predescu, S.M. Vogel, A.B. Malik, Functional and morphological studies of protein transcytosis in continuous epithelia. Am. J. Physiol. 287, L895–901 (2004).

70. V. Triacca, E. Guc, W.W. Kilarski, M. Pisano, M. Swartz M, Transcellular pathways in lymphatic endothelial cells regulate changes in solute transport by fluid stress. Circulation Research 120,1440–1452 (2017).

71. M. Van Lessen, S. Shibata-Germanos, A. van Impel, T.A. Hawkins, J. Rihel, S. Schulte-Merker, Intracellular uptake of macromolecules by brain lymphatic endothelial cells during zebrafish embryonic development. eLife 6, e25932 (2017).

72. R.G. Parton, K. Simons, The multiple faces of caveolae. Nat. Rev. Mol. Cell Biology 8, 185–194 (2007).

73. C. LeRoy, J.L. Wrana, Clathrin and non-clathrin mediated endocytotic regulation of cell signaling. Nat. Rev. Mol. Cell Biology 6, 112–126 (2005).

74. K.H. Andres, M. von During, K. Muszynski, R.F. Schmidt, Nerve fibres and their terminals of the dura mater encephali of the rat. Anat Embryol 175, 289–301 (1987).

75. M. Krohn, C. Lange, J. Hofrichter, K. Scheffier, J. Stenzel, J. Steffen, T. Schumacher, T. Bruning, A.S. Plath, F. Alfen, A. Schmidt, F. Winter, K. Rateitschak, A. Wree, J. Gsponer, L.C. Walker, J. Pahnke, Cerebral amyloid-β proteostasis is regulated by the membrane transport protein ABCC1 in mice. JCI 121(10), 3924–31 (2011).

76. M. Krohn, A. Bracke, Y. Avchalumov, T. Schumacher, J. Hofrichter, K. Paarmann, C. Frohlich, C. Lange, T. Bruning, O. von Bohien und Halbach, J. Pahnke, Accumulation of murine amyloid-β mimics early Alzheimer’s disease. Brain 138(8), 2370-82 (2015)

77. W.B. Stine, L. Jungbauer, C. Yu, M.J. LaDu, Preparing synthetic Aβ in different aggregation states. Methods Mol. Biol. 670, 13–32 (2011).

78. J.R. Cirrito, R. Deane, A.M. Fagan, M.L. Spinner, M. Parsadian, M.B. Finn, H. Jiang, J.L. Prior, A. Sagare, K.R. Bales, S.M. Paul, B.V. Zlokovic, D. Piwica-Worms, D M. Holtzman, P-glycoprotein (ABCB1) deficieny at the blood-brain barrier increased amyloid-β deposition in an Alzheimer disease mouse model. J. Clin. Invest. 115, 3285–3290 (2005).

79. A. Brenn, M. Grube, M. Peters, A. Fischer, G. Jedlitschky, H.K. Kroemer, R.W. Warzok, S. Vogelgesang, Beta-amyloid downregulates ABCB1 expression at the blood-brain barrier in mice, Int. J. Alz. Dis., doi:10.4061/2011/690121

80. J. Pahnke, O. Wolkenhauer, M. Krohn, L.C. Walker, Clinico-pathologic function of cerebral ABC transporters-implications for the pathogenensis of Alzheimer’s disease. Curr. Alz. Res. 5(4), 396–405 (2008).

81. C. Janson, U.S. patent application, “Lymphatic & arachnoid cells for CNS discovery” (2017).

82. M. Foldi, E. Foldi, R.H.K. Strosenreuther, S. Kubik, Eds., Foldi’s Textbook of Lymphology (Elsevier, Munchen, 2011), p.4–5.

83. S. Podgrabinska, P. Braun, P. Velasco, B. Kloos, M.S. Pepper, D.G. Jackson, M. Skobe, Molecular characterization of lymphatic endothelial cells. PNAS 99(25), 16069–16074 (2002).

84. N. Wick, P. Saharinen, J. Saharinen, E. Gurnhofer, C.W. Steiner, I. Raab, D. Stokic, P. Giovanoli, S. Buchsbaum, A. Burchard, S. Thurner, K. Alitalo, D. Kerjaschki, Transcriptional comparison of human dermal lymphatic endothelial cells ex vivo and in vitro. Physiol. Genomics 28, 179–192 (2007).

85. J.C. Lambert et al., Meta-analysis of 74,046 individuals identifies 11 new susceptibility loci for Alzheimer’s disease. Nat. Genetics 45(12), 1452–1458 (2013).

86. H. W. Cushing, Studies on the cerebrospinal fluid. Journal of Medical Research 31(1), 1–19 (1914).

87. J.E. Schnitzer, P. Oh, E. Pinney, J. Allard, Filipin-sensitive caveolae-mediated transport in endothelium: reduced transcytosis, scavenger endocytosis, and capillary permeability of select macromolecules. J. Cell Biol. 127(5), 1217–32 (1994).

88. D. Predescu, S.M. Vogel, A.B. Malik, Functional and morphological studies of protrein transcytosis in continuous endothelia. Am. J. Physiol. Lung Cell. Mol. Physiol. 287, L895–901 (2004).

89. S. Pavlides, A. Tsirigos, I. Vera, N. Flomenberg, P.G. Frank, M.C. Casimiro, C. Wang, P. Fortina, S. Addya, R.G. Pestell, U.E. Martinez-Outschoorn, F. Sotgia, M.P. Lisanti, Loss of stromal caveolin-1 leads to oxidative stress, mimics hypoxia and drives inflammation in the tumor microenvironment, conferring the “reverse Warburg effect”: a transcriptional informatics analysis with validation. Cell Cycle 9(11), 2201–19 (2010).

90. B.P. Head, J.N. Peart, M. Panneerselvam, T. Yokoyama, M.L. Pearn, I.R. Niesman, J.A. Bonds, J.M. Schilling, A. Miyanohara, J. Headrick, S.S. Ali, D.M. Roth, P.M. Patel, H.H. Patel, Loss of caveolin-1 accelerates neurodegeneration and aging. PLoS One. 5(12), e15697 (2010).

91. S.B. Gaudreault, D. Dea, J. Poirier, Increased caveolin-1 expression in Alzheimer’s disease brain. Neurobiol. Aging 25(6),753–9 (2004).

92. R.S. Thomas, M.J. Lelos, M.A. Good, E.J. Kidd, Clathrin-mediated endocytic proteins are upregulated in the cortex of the Tg2576 mouse model of Alzheimer’s disease-like amyloid pathology. Biochem Biophys Res Commun. 415(4), 656–61 (2011).

93. J.P. Hoffmann, Generalized linear models : an applied approach (Boston, Pearson, 2004).

94. S. Greenland, Estimating standardized parameters from generalized linear models. Stat Med, 10(7), 1069– 1074 (1991).

95. Y. Benjamini, Y. Hochberg, Controlling the False Discovery Rate: A Practical and Powerful Approach to Multiple Testing. Journal of the Royal Statistical Society. Series B (Methodological), 57(1), 289–300 (1995).

96. E. Jung, D. Gardner, D. Choi, E. Park, Y.S. Seong, S. Yang, J. Castorena-Gonzalez, A. Louveau, Z. Zhou, G.K. Lee, D.P. Perrault, S. Lee, M. Johnson, G. Daghlian, M. Lee, Y.J. Hong, Y. Kato, J. Kipnis, M.J. Davis, A.K. Wong, Y.K. Hong, Development and characterization of a novel Prox1-eGFP lymphatic and Schlemm’s canal reporter rat. Sci. Rep. 7, 5577 (2017).

97. S. Tsutsumi, M. Nakamura, T. Tabuchi, Y. Yasumoto, M. Ito, Cranial arachnoid protrusions and contiguous diploic veins in CSF drainage. Am. J. Neuroradiol. 35(9), 1735–9 (2014).

98. S. Tsutsumi, H. Ono, Y. Yasumoto, Pile driving into the skull and suspending the bridging veins? An undescribed role of arachnoid granulations. Surg. Rad. Anatomy 39(5), 541–545 (2017).

